# Auditory tracking magnitude, temporal alignment, and anatomical distribution during wakefulness and propofol-based anesthesia

**DOI:** 10.64898/2026.10.23.761298

**Authors:** Hao Zhu, Chen Yao, Zhili Han, Wenfang Zhou, Yuanqing Wang, Sixian Li, Patrick C M Wong, Xing Tian, Xiangbin Teng

**Affiliations:** Brain and Mind Institute (BMI), The Chinese University of Hong Kong, Shatin, Hong Kong SAR, China; Department of Psychology, The Chinese University of Hong Kong, Shatin, Hong Kong SAR, China; NYU-ECNU Institute of Brain and Cognitive Science at NYU Shanghai, Shanghai, China; Department of Linguistics and Modern Languages, The Chinese University of Hong Kong, Shatin, Hong Kong SAR, China; Department of Neurosurgery, The National Key Clinic Specialty, Shenzhen Key Laboratory of Neurosurgery, the First Affiliated Hospital of Shenzhen University, Shenzhen Second People’s Hospital, Shenzhen, China; NingboTech University, Ningbo, China; Division of Arts and Sciences, New York University Shanghai, Shanghai, China; Department of Otolaryngology, Shenzhen Luohu People’s Hospital, Shenzhen, China

## Abstract

General anesthesia attenuates auditory responses, but response magnitude alone does not describe how residual activity aligns with sound over time or varies across the brain. We jointly characterized these aspects of auditory tracking using stereoelectroencephalography (sEEG) in 13 patients with drug-resistant epilepsy during wakefulness and clinically defined unresponsiveness under propofol-based anesthesia. The same rhythmic humming sequences were presented at approximately 1.61 Hz in both sessions. During anesthesia, stimulus-rate power showed widespread reductions and phase alignment with the stimulus rhythm was less consistent, although responses remained detectable at some recording channels. During wakefulness, response strength decreased with cortical surface distance from posteromedial Heschl’s gyrus; this distance-response relationship was weaker during anesthesia. In a contrastive learning analysis (CEBRA), low-dimensional embeddings derived from anesthesia recordings yielded lower acoustic-envelope decoding accuracy on held-out data and lower consistency across model fits. Together, these findings characterize how attenuated auditory tracking is expressed jointly in response magnitude, temporal alignment, and anatomical distribution.

## Introduction

General anesthesia produces clinically defined unresponsiveness and widespread changes in brain activity and functional connectivity^1–3^, while its effects on auditory processing vary across regions and response types. Human functional magnetic resonance imaging (fMRI) studies show that auditory cortical responses can remain detectable during sedation and anesthesia even as responses associated with speech comprehension or stimulus category are attenuated^4,5^. In macaques, responses to auditory inputs can persist while local and global novelty change^6^. Scalp EEG recordings show reduced auditory steady-state responses during propofol sedation^7^ and general anesthesia with other agents^8^. Human intracranial studies further demonstrate differences across cortical regions, response components, and levels of auditory sequence processing^9–11^. These findings establish that auditory responsiveness under anesthesia is heterogeneous and motivate examining how residual tracking is expressed across time and recording locations.

Auditory processing also varies with attentional engagement and arousal outside anesthesia. Research on auditory event-related potentials (ERPs) distinguishes responses that can occur without directed attention from those that depend more strongly on attentional engagement^12,13^. During wakefulness, tracking of unattended speech, syllable-rate tracking during distraction, and tracking of repeated artificial words demonstrate that some auditory regularities are represented when attention is directed elsewhere^14–16^. During sleep, substantial auditory spiking and high-gamma responses can coexist with reduced alpha-beta desynchronization^17^, and local slow waves modulate temporal-lobe evoked responses^18^. These observations place anesthesia within a broader question of how a listener’s state shapes the strength and timing of sensory responses.

Rhythmic stimulation provides a common acoustic reference for examining these aspects of auditory processing. Studies relate response phase to detection performance^19^ and show phase-sensitive tracking across timescales^20^, within broader accounts of entrainment and selective listening^21,22^. Measuring activity relative to a recurring stimulus therefore allows both response magnitude and temporal alignment to be described. Such measurements characterize tracking without, on their own, establishing whether it arises from repeated evoked responses, endogenous oscillatory alignment, or a combination of processes^23^.

The present study extends previous regional and temporal characterizations of auditory responses under anesthesia^9–11^ by examining response magnitude, temporal alignment, and anatomical distribution relative to a common stimulus rhythm. Stimulus-rate power describes the strength of tracking, phase measures describe its temporal alignment, and anatomical analyses describe how these responses vary across recording sites. Evaluating these measures in the same participants under the same stimulation conditions characterizes how attenuation is expressed across these aspects of the response. The measurements provide complementary descriptions instead of assuming that their state differences arise from independent mechanisms.

Here, we compared stereoelectroencephalography (sEEG) recordings from 13 patients with drug-resistant epilepsy during wakefulness and propofol-based anesthesia (Fig. 1). The same sequences of humming sounds were presented in both sessions, with individual sounds recurring at approximately 1.61 Hz; awake participants were instructed to listen passively. We quantified stimulus-rate power and phase consistency and examined their anatomical distribution, including the relationship between response strength and cortical surface distance from posteromedial Heschl’s gyrus. As a complementary analysis, we used CEBRA to learn low-dimensional embeddings conditioned on the acoustic envelope^24^ and evaluated envelope decoding on held-out data and consistency across repeated model fits. This within-participant comparison provides a joint, stimulus-referenced account of auditory tracking across the recorded clinical states.

**Fig. 1.**
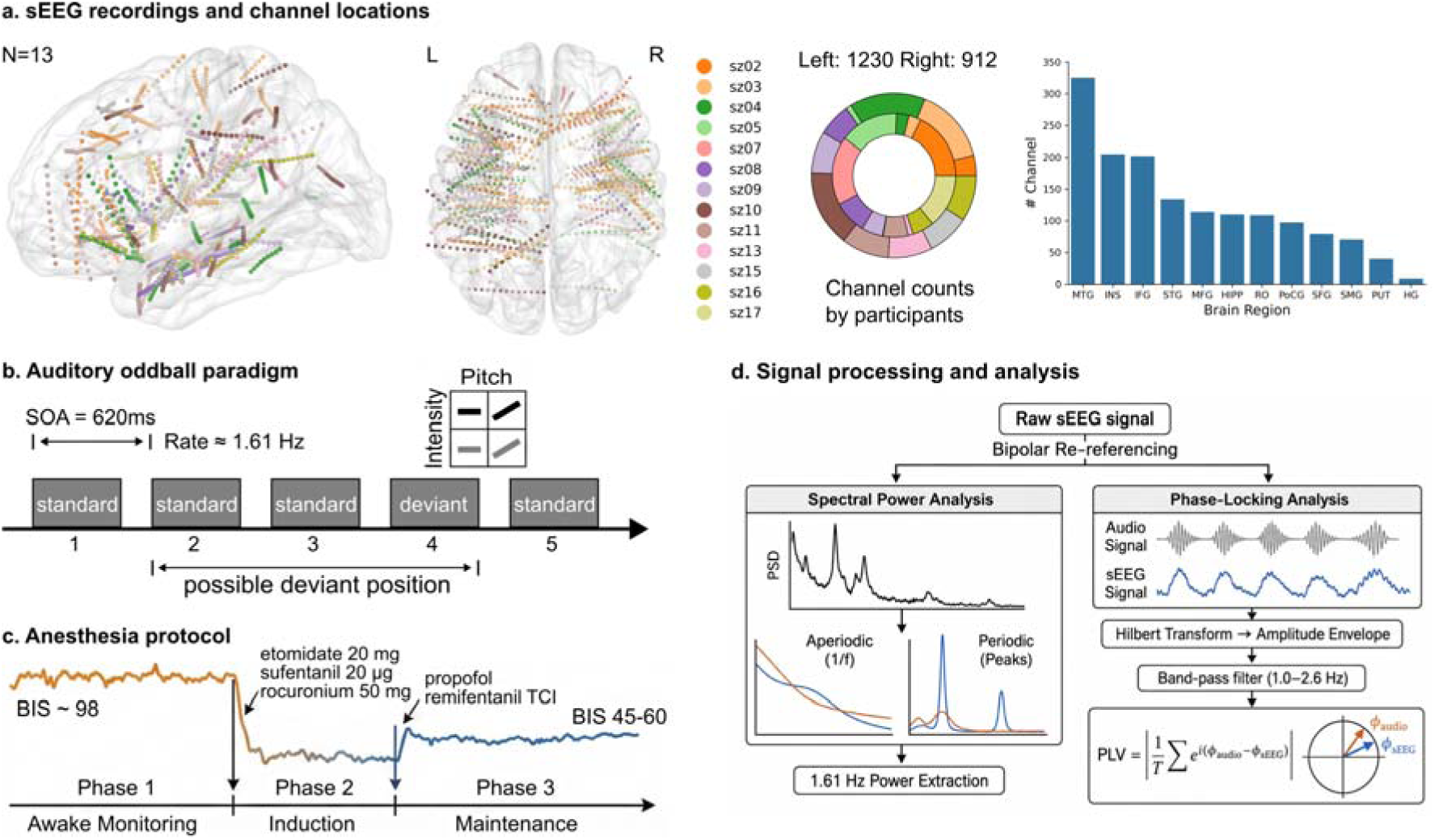
Experimental design and analytical framework. (a) Anatomical distribution of sEEG depth electrodes. A total of 2,142 contacts (1,230 left, 912 right) from 13 participants were localized and normalized to the MNI standard space. The bar chart (right) displays the channel count distribution across recorded brain regions. (b) Rhythmic auditory stimulation paradigm. Participants heard five-sound sequences of humming sounds with a stimulus onset asynchrony (SOA) of 620 ms (approximately 1.61 Hz). Each sequence contained four standard sounds and one deviant differing in pitch contour (flat versus rising) or intensity (60 versus 80 dB SPL). Standard and deviant sounds were pooled for the present rhythmic-tracking analyses. (c) Schematic anesthesia protocol. Anesthesia recordings were obtained during implantation surgery; wakefulness recordings were obtained later during ward monitoring. (d) Signal processing pipeline. Raw sEEG signals were re-referenced using a bipolar montage. Spectral power analysis utilized the FOOOF algorithm^48^ to isolate periodic peaks at the stimulus frequency (1.61 Hz) from the aperiodic background. PLV and mean phase angles at stimulus onset were computed by extracting the instantaneous phase of the signal envelope via the Hilbert transform, band-pass filtered at the stimulus rate (1.0-2.6 Hz).

## Results

### Lower stimulus-rate power during anesthesia

To compare auditory tracking magnitude between wakefulness and propofol-based anesthesia, we measured spectral power at the stimulus rate (1.61 Hz; 620 ms SOA) as participants listened to five-item sequences of humming sounds. Full details of stimulus construction, sequence structure, and block design are provided in the Methods (Fig. 1).

During wakefulness, auditory stimuli elicited robust oscillatory responses across a distributed network of cortical and subcortical regions (Fig. 2a; see Supplementary Fig. 5 for an illustration of the full spectral and phase pipeline applied to a representative right-STG channel). The distribution of auditory-responsive channels across participants and ROIs in each state is reported in Supplementary Table 2. FOOOF spectral parameterization yielded high-quality fits across all recordings (mean R² = 0.982, SD = 0.023, median = 0.987; Supplementary Table 3). Only 25 of 3,050 channel-level fits (0.8%) fell below R² = 0.90. Overall FOOOF fit quality was high in both conditions (mean R²: anesthesia, 0.984; wakefulness, 0.980). During the anesthesia session, stimulus-rate power was lower than during wakefulness. A comparison of the aperiodic-adjusted power distributions using a linear mixed-effects model with participant as a random intercept (see Methods) revealed that neural responses were significantly stronger during wakefulness compared to anesthesia (β = 5.07 dB, z = 19.01, p < 0.001, Fig. 2d). This suppression was widespread, spanning the auditory pathway as well as frontal and motor regions. ROI analysis showed lower power during anesthesia in primary auditory areas (Heschl’s Gyrus, HG), secondary auditory areas (Superior Temporal Gyrus, STG; Middle Temporal Gyrus, MTG), and association sensorimotor cortices including the Rolandic Operculum (RO), Inferior Frontal Gyrus (IFG), Postcentral Gyrus (PoCG), and Fusiform Gyrus (FG) (all p < 0.01, Fig. 2c). Lower power was also observed in the hippocampus (HIPP), insula (INS), and putamen (PUT). This widespread attenuation is consistent with prior reports of reduced auditory cortical responses under propofol, both in scalp recordings^7^ and intracranial studies showing disruption beyond primary auditory cortex^9^.

**Fig. 2.**
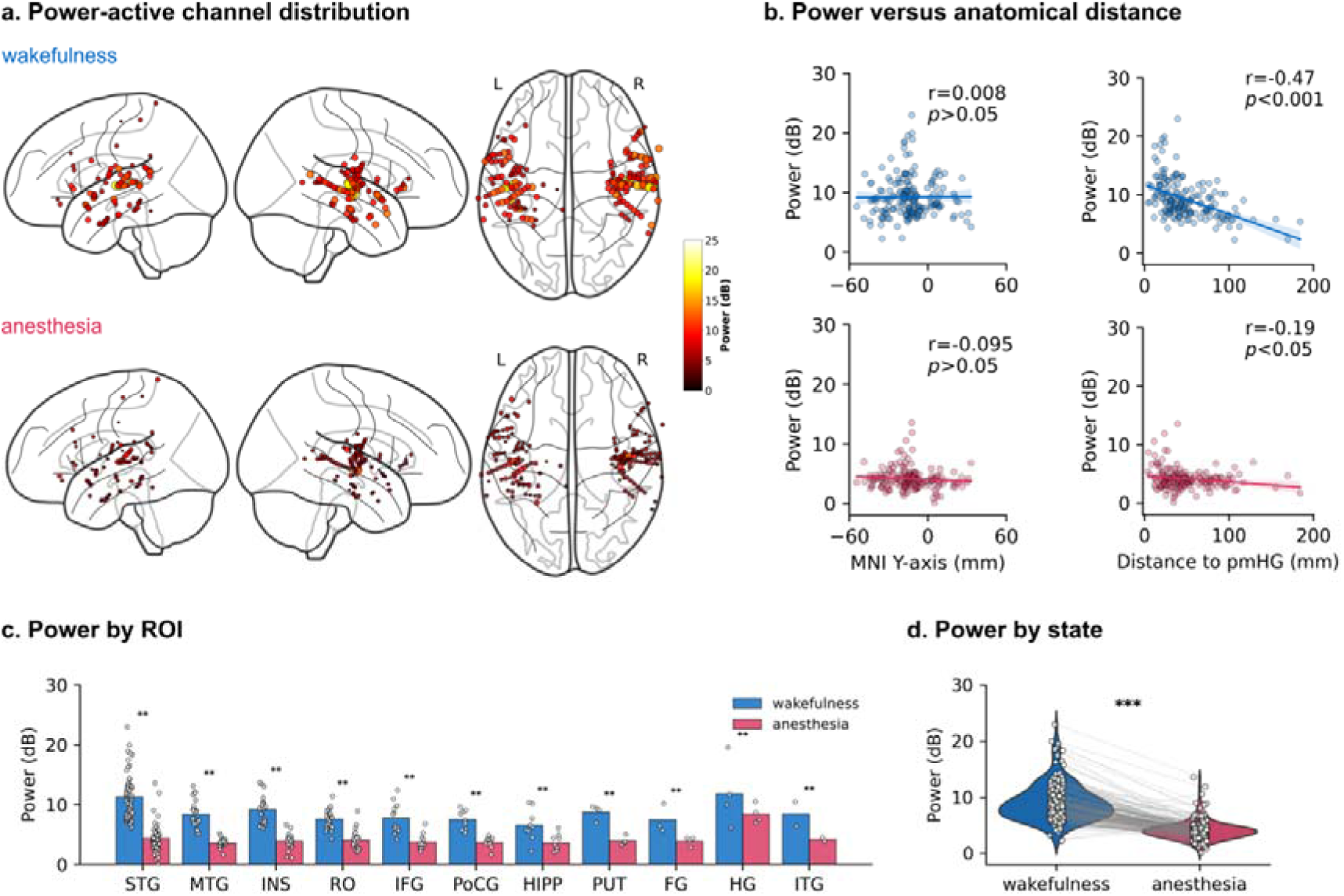
Stimulus-rate power and its spatial distribution during wakefulness and anesthesia. (a) Spatial distribution of auditory tracking power across the brain during wakefulness (top) and anesthesia (bottom). Active channels are projected onto a glass brain in MNI space, with color and size indicating the magnitude of aperiodic-adjusted power at 1.61 Hz. (b) Spatial gradient (distance-response) analysis. Scatter plots correlate spectral power (y-axis) with the MNI Y-coordinate (anterior-posterior axis, left column) and the geodesic distance from the pmHG along the cortical surface (right column). Solid lines indicate the best-fit linear regression, and shaded bands show the 95% confidence intervals around the regression line. Power was negatively associated with geodesic distance from pmHG during wakefulness (r = −0.47, p < 0.001), with a weaker association during anesthesia (r = −0.19, p = 0.011). Anterior-posterior correlations were small (wakefulness: r = 0.008, p = 0.914; anesthesia: r = −0.095, p = 0.209). (c) ROI analysis. Bar charts show mean power across anatomical regions. Power was lower during anesthesia in primary (HG) and secondary (STG, MTG) auditory cortices, as well as association areas (RO, IFG) and subcortical structures (HIPP, PUT) (**: p < 0.01, channel-stratified permutation test). (d) Global state comparison. Violin plots illustrate the distribution of power across all responsive channels, showing a significant global attenuation during anesthesia compared to wakefulness (***: p < 0.001).

To determine whether anesthesia-responsive channels constitute a subset of the wakefulness-responsive population or represent a distinct set, we characterized the overlap between the two sets of significant channels. Of 171 power-responsive channels during wakefulness and 24 during anesthesia, 17 appeared in both states (Jaccard similarity = 9.6%). Most power-responsive channels under anesthesia (70.8%, 17/24) were also responsive during wakefulness, indicating that residual auditory tracking during anesthesia largely persists within the same channels that are active during wakefulness rather than emerging in a new population. All power-responsive channels under anesthesia in STG (10/10), MTG (1/1), and INS (2/2) were also responsive during wakefulness. In contrast, a small number of channels in IFG and HIPP became significant only under anesthesia, and several ROIs (PoCG, PUT, FG) lost all responsive channels entirely (Supplementary Table 2).

### Spatial distribution of stimulus-rate power during wakefulness and anesthesia

Auditory tracking at the 1.61 Hz stimulus presentation rate remained detectable during anesthesia despite the overall reduction in stimulus-rate power (Fig. 2a). We next examined how the spatial distribution of response strength differed between wakefulness and anesthesia. We quantified the distance-response relationship, here termed the *spatial gradient*, using geodesic distance along the cortical surface from posteromedial Heschl’s gyrus (pmHG, TE1.1), used as an anatomical reference for auditory core regions (see Methods). This analysis assessed how stimulus-rate power varied across recording sites relative to this reference.

During wakefulness, channels farther from pmHG tended to have lower stimulus-rate power (Pearson’s r = −0.47, p < 0.001; Fig. 2b). The association was weaker during anesthesia (r = −0.19, p = 0.011). By comparison, correlations with anterior-posterior position were small in both states (wakefulness: r = 0.008, p = 0.914; anesthesia: r = −0.095, p = 0.209). These results describe a distance-response association that was more pronounced during wakefulness. The participant-fixed-intercept analysis showed an anesthesia-minus-wakefulness difference in power–distance slope of +3.24 dB/100 mm (95% CI: [1.14, 5.31], p = 0.017; Supplementary Fig. 6c).

We also examined how rescaling response values affected the distance-response relationship in a simulation (Supplementary Fig. 6c). Uniform scaling by factors from 1.0 to 0.1 left the correlation unchanged, consistent with the identity r(X, aY) = r(X, Y) for a > 0. In contrast, the simulated distance-dependent attenuation progressively steepened the negative gradient. Thus, uniform rescaling alone did not reproduce the weaker association observed during anesthesia, although this simulation does not exclude effects of noise, response detectability, or other forms of attenuation.

### Phase consistency during wakefulness and anesthesia

We next examined the timing of auditory responses relative to the stimulus rhythm^20,22,25,26^. For each channel, we quantified onset phase concentration using the mean resultant length (R) of the filtered sEEG envelope phase at successive tone onsets in the trial-averaged response^27^. Larger R values indicate greater concentration of onset phases. We also measured the phase-locking value (PLV) between the averaged acoustic and sEEG envelopes to characterize their phase relationship over time. Onset phase concentration was lower during anesthesia than during wakefulness (wakefulness: R = 0.557 ± 0.022; anesthesia: R = 0.233 ± 0.011; mean ± s.e.m.; Fig. 3c). The paired state comparison was significant (Wilcoxon signed-rank test, W = 1138, p < 0.001). Envelope PLV showed the same direction of change (Supplementary Fig. 1). Together, these results indicate reduced consistency of stimulus-related phase alignment during anesthesia.

**Fig. 3.**
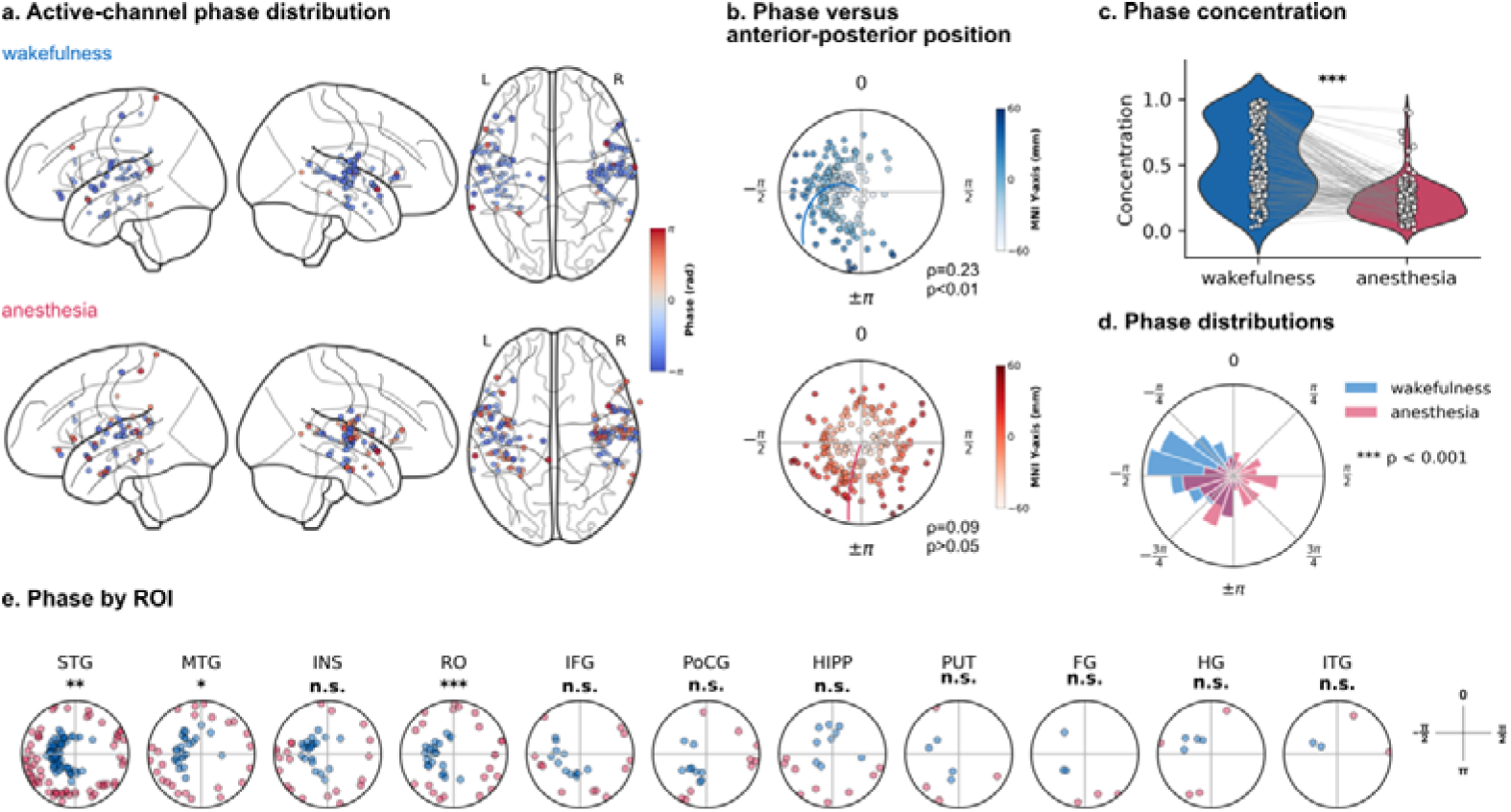
Phase measures of auditory tracking during wakefulness and anesthesia. (a) Topography of neural phase angles. Glass brain visualization of the mean phase angle for responsive channels during wakefulness (top) and anesthesia (bottom). (b) Circular-linear correlation analysis. Plots depict the relationship between the phase angle of neural entrainment and the anterior-posterior anatomical axis (MNI Y-coordinate). The association was stronger during wakefulness (ρ = 0.23, p < 0.01) and weaker during anesthesia (ρ = 0.09, p > 0.05). (c) Onset phase concentration Violin plots show the mean resultant length (R) of sEEG envelope phase at tone onsets in each channel’s trial-averaged response. Larger values indicate greater concentration of onset phases. (d) Polar histograms of phase distribution across all channels. Angles are expressed in radians. Reported mean angles use the interval [0, 2π], while polar tick labels show the equivalent angles wrapped to [-π, π]. The phase angle distribution was more concentrated during wakefulness (mean angle = 4.73 rad) than during anesthesia; the distribution differed between states (Watson’s U^2^ = 1.575). (e) ROI-specific phase analysis. Circular plots show phase distributions for key anatomical regions. Differences in phase angle distributions were observed in the STG, MTG, and RO (***: p < 0.001; **: p < 0.01; *: p < 0.05; n.s.: not significant).

We used simulations to examine how spectral power and envelope PLV responded to manipulations of response amplitude and phase alignment (Supplementary Fig. 6a). Changing amplitude altered power by 27.75 dB and PLV by 0.020, equivalent to 2.4% of the PLV effect produced by the phase manipulation. Changing phase alignment altered PLV by 0.831 and power by 0.32 dB, equivalent to 1.1% of the power effect produced by the amplitude manipulation. These results show that each measure was more sensitive to its corresponding manipulation within the tested signal model and parameter range. However, methodological work on intertrial phase coherence shows that changes in power and evoked responses can alter measured phase consistency without a change in the underlying phase^28^. The finding identifies a limitation when interpreting phase estimates from noisy recordings. The simulations therefore support different sensitivities of the measures under the modeled conditions, but do not establish their independence in the recorded data.

We also examined the distribution of mean phase angles at tone onsets across the analyzed channels (Fig. 3a, d). The distribution was more concentrated during wakefulness (mean angle = 4.73 rad, κ = 1.61) than during anesthesia (mean angle = 3.40 rad, κ = 0.40). The distributions differed between states (Watson’s *U*^2^= 1.575, critical value = 0.187 at α = 0.05). Circular–linear analysis (Fig. 3b) showed an association between mean phase angle and anterior-posterior position during wakefulness (ρ = 0.228, p < 0.01) and a weaker association during anesthesia (ρ = 0.086, p = 0.515). Related PLV spatial analyses are shown in Supplementary Fig. 1.

We compared phase-angle distributions between wakefulness and anesthesia across eleven anatomical ROIs using the nonparametric Wheeler-Watson test^29^ (Fig. 3e). State differences were observed in auditory regions, including STG (W = 15.83, p < 0.01), MTG (W = 13.47, p < 0.05), and RO (W = 19.96, p < 0.001). These results identify state-related differences in phase-angle distributions across sampled cortical regions.

### Acoustic-envelope decoding from CEBRA embeddings

To complement the spectral, phase, and spatial analyses, we examined auditory tracking using a low-dimensional representation of the multichannel sEEG recordings. This approach draws on neural manifold frameworks that summarize high-dimensional activity as trajectories in a latent space^30–32^. We used CEBRA^24^ to learn three-dimensional embeddings conditioned on the acoustic envelope (Fig. 4a). We assessed these representations by decoding the envelope from held-out data and measuring embedding consistency across model fits. Representative embeddings from wakefulness and anesthesia are shown in Fig. 4b.

**Fig. 4.**
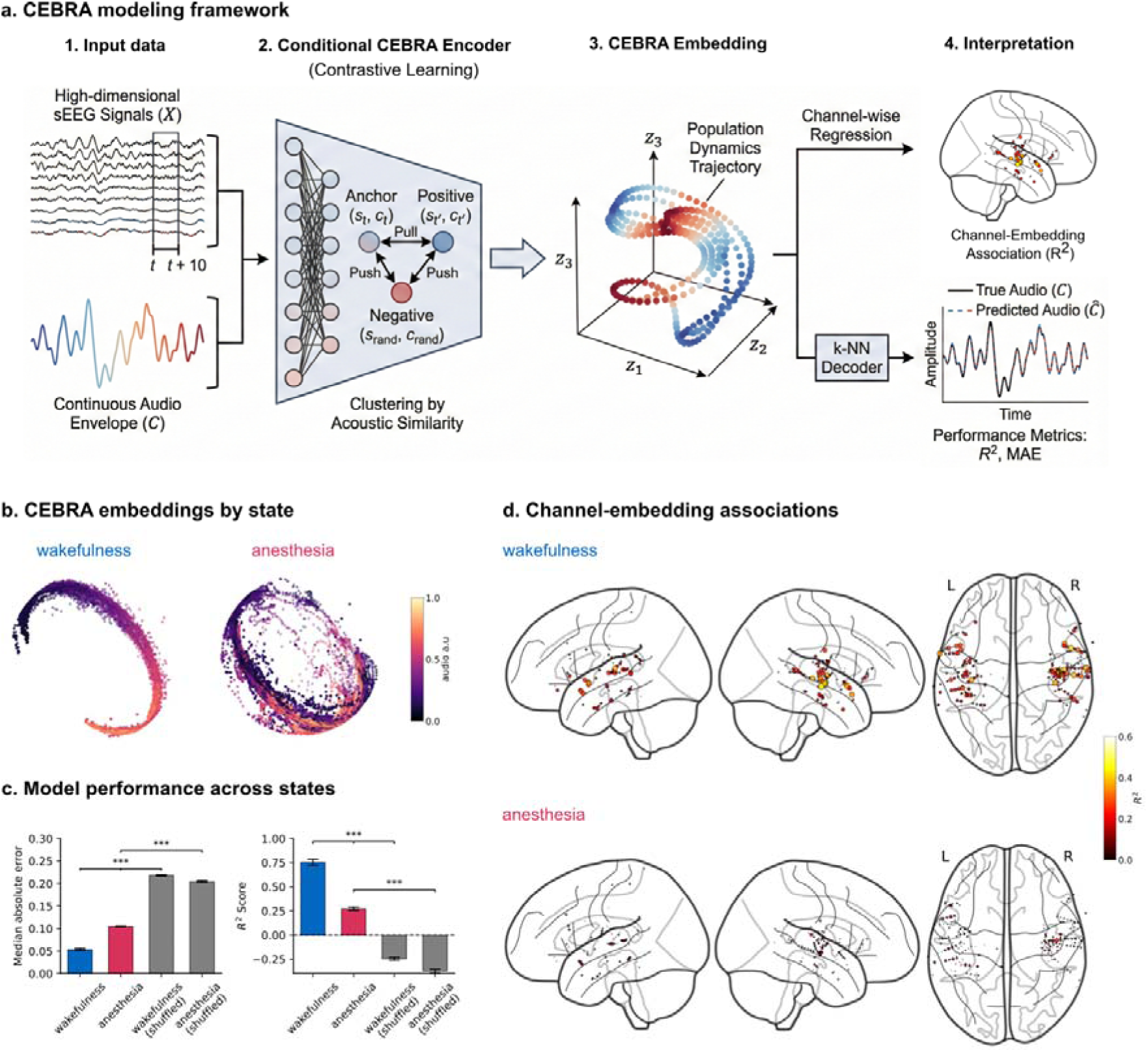
CEBRA embeddings and acoustic-envelope decoding during wakefulness and anesthesia. (a) The CEBRA modeling framework. The CEBRA encoder mapped high-dimensional sEEG signals to a latent representation, using the acoustic envelope as a conditioning variable during training. (b) Representative three-dimensional CEBRA embeddings during wakefulness (left) and anesthesia (right). These plots illustrate model-derived representations of the sEEG data. (c) Acoustic-envelope decoding performance. Decoding performance (left: MAE; right: R^2^) of the audio envelope from neural embeddings. Bars show means across model fits per condition, and error bars show s.e.m. R² significance annotations correspond to two-sided Wilcoxon signed-rank comparisons with FDR correction (***, p < 0.001). The true model in wakefulness significantly outperforms both the anesthesia model and shuffled controls. (d) Channel-embedding associations. Brain maps show R² values from separate regressions relating each channel’s sEEG activity to the embedding coordinates, averaged across model runs.

Held-out acoustic-envelope decoding was more accurate during wakefulness ( = 0.753±0.032, MAE = 0.054±0.003) than during anesthesia ( = 0.272±0.017, MAE = 0.105±0.001; mean±s.e.m.; Fig. 4c). We compared models trained with the original envelope labels against controls trained with shuffled labels (see Methods). During wakefulness, decoding was higher with the original labels than with shuffled labels (0.753 versus −0.246, p < 0.001), and MAE was lower (p < 0.001). Anesthesia decoding also exceeded its shuffled baseline, despite the lower performance relative to wakefulness.

Embedding consistency was lower during anesthesia. In the consistency analysis, scores were 0.561 during wakefulness and 0.191 during anesthesia (Supplementary Fig. 2b; p < 0.001). The wakefulness score also exceeded that of the shuffled controls (0.561 versus 0.135, p < 0.001). These results describe differences in the reproducibility of the learned embeddings across model fits.

Separate regressions relating each channel’s activity to the embedding coordinates showed stronger associations in HG, STG, and MTG during wakefulness than during anesthesia (Fig. 4d). Broadband sEEG amplitude at the model input was similar across states (anesthesia-to-wakefulness ratio = 1.02). We next compared training and validation performance in each state (Supplementary Fig. 3). Training mean goodness-of-fit (GoF) was 1.05 bits during wakefulness and 1.04 bits during anesthesia. The training-validation GoF gap was larger during anesthesia (0.977 ± 0.009) than during wakefulness (0.435 ± 0.031; Wilcoxon signed-rank test, p < 0.001). Decoding R² showed the same pattern, with gaps of 0.722 ± 0.017 during anesthesia and 0.242 ± 0.032 during wakefulness.

Finally, we assessed validation performance after training models with 20%, 40%, 60%, 80%, and 100% of the available training epochs (Supplementary Fig. 4). Wakefulness validation performance remained within a similar range across these fractions (R² = 0.80-0.84). Anesthesia validation performance did not improve as the training fraction increased, while the training-validation gap widened. Across the evaluated training fractions and data partitions, anesthesia models retained lower validation performance within the available dataset and tested training settings.

## Discussion

Our findings characterize the temporal alignment and anatomical distribution of attenuated auditory tracking under propofol-based anesthesia. Using the same auditory sequences in the same participants, we observed lower stimulus-rate power alongside less consistent phase alignment and a weaker relationship between response strength and distance from posteromedial Heschl’s gyrus. This joint characterization extends prior descriptions of regional and temporal differences in auditory responses under anesthesia^9–11^ by relating response magnitude, timing, and anatomical location to a common stimulus rhythm. Complementary CEBRA analyses showed lower held-out acoustic-envelope decoding and lower embedding consistency across model fits. Together, the measurements describe how auditory tracking differs across the recorded states, while leaving open the extent to which their concurrent changes share a dependence on stimulus-related signal strength.

Residual auditory responses remained detectable during anesthesia despite the overall reduction in stimulus-rate power. Most channels with detectable power responses under anesthesia also responded during wakefulness (17/24, 70.8%; Supplementary Table 2), indicating that residual tracking largely involved sites responsive in both states. A small number of IFG and HIPP channels met the power-responsiveness criterion only during anesthesia. Because the criterion was applied separately in each state, these threshold crossings do not by themselves show that anesthesia recruited new auditory responses. This pattern is consistent with neuroimaging evidence of residual auditory responsiveness under anesthesia^4–6^ and intracranial findings of uneven anesthetic effects across regions and response components^9–11^. Sleep provides a related example in which substantial auditory responses persist alongside altered feedback signaling and modulation by slow waves^17,18^. These findings make the timing and anatomical distribution of detectable responses informative alongside their overall magnitude.

Lower onset phase concentration and envelope PLV during anesthesia indicated less consistent alignment with the 1.61 Hz stimulus rhythm. Phase-angle distributions also differed between states, and their association with anterior-posterior position was weaker during anesthesia. These observations describe changes in both within-channel phase consistency and the distribution of response timing across recording sites. Previous studies relate response phase to perceived event timing^25^ and to speech structure beyond low-level acoustic fluctuations^26^, within broader accounts of entrainment and auditory attention^21,22^. In the present non-speech paradigm, however, phase locking could arise from repeated evoked responses, endogenous oscillatory alignment, or both^21^. The measurements therefore characterize temporal alignment without establishing its generating mechanism or its consequences for perception.

The simulations support using power and phase measures together while defining a limit on their interpretation. Within the tested signal model and parameter range, power and envelope PLV were preferentially sensitive to amplitude and phase manipulations, respectively (Supplementary Fig. 6a). Auditory MEG findings that phase-based classification can outperform power-based classification provide a related rationale for considering both measures^20^. Nevertheless, methodological work shows that power and evoked-response changes can affect intertrial phase-coherence estimates^28^. Although that measure differs from the envelope PLV used here, stimulus-related signal strength remains a possible contributor to the observed phase differences. The simulations do not establish power-phase independence in the recordings.

The distance-response analysis adds an anatomical description of the attenuation. During wakefulness, stimulus-rate power decreased with geodesic distance from pmHG; this relationship was weaker during anesthesia, whereas anterior-posterior power correlations were small in both states (Fig. 2). Response strength was therefore more closely associated with proximity to the auditory-core reference during wakefulness. In the tested simulations, uniform positive rescaling preserved the correlation and distance-dependent attenuation steepened the negative gradient (Supplementary Fig. 6c). Noise, response detectability, and channel selection can also affect this relationship, so the result identifies a difference in the distribution of response strength across sampled sites instead of establishing a specific anatomical mechanism.

CEBRA provided a complementary view of stimulus-related information in multichannel activity. Population representations summarize relationships across distributed activity patterns ^30,33^; related intracranial work has characterized auditory resting-network geometry^34^ and cortical dynamics associated with word forms^35^. Here, CEBRA used the acoustic envelope to guide representation learning^24^, allowing us to ask how accurately that envelope could be recovered from held-out recordings. Decoding accuracy was lower during anesthesia but remained above the shuffled baseline (Fig. 4c), indicating that the fitted model recovered residual envelope information with reduced accuracy. Stronger channel-embedding associations in HG, STG, and MTG during wakefulness further connected the representation to activity in these sampled auditory regions (Fig. 4d); these marginal associations do not identify unique channel contributions. The selected three-dimensional embedding provides a compact description of activity that may contain higher-dimensional structure^31^.

Mean embedding consistency was lower during anesthesia across repeated model fits (Supplementary Fig. 2), and decoding performance remained lower across the tested training fractions (Supplementary Fig. 4). Broadband input amplitude was similar between states (anesthesia-to-wakefulness ratio = 1.02), although the stimulus-related component could still differ. These results describe the performance and reproducibility of the fitted representations under the tested conditions and complement broader work on anesthetic changes in cortical stability and network dynamics^36–38^. CEBRA guides learning with the supplied stimulus variable, whereas PCA summarizes dominant variance in the recordings. A direct comparison with PCA was not performed, so these analyses do not establish that CEBRA is preferable to other representations. Their role here is to supplement the power, phase, and anatomical measurements with stimulus-referenced visualization and decoding.

Attention remains a possible contributor to the state differences, but two considerations temper an interpretation based solely on its withdrawal. First, auditory tracking is not uniformly contingent on directed attention: cortical responses track unattended speech^14^, syllable-rate tracking persists during auditory or visual distraction^15^, and repeated artificial words can be tracked during a demanding visual task^16^. These findings show that reduced attentional engagement need not abolish low-level tracking. Second, the present analyses measured tracking of the acoustic stimulus rhythm during passive listening and pooled across sound types; participants were not required to identify sequences or make an overt response. The comparison therefore does not depend on successful performance of an active auditory task. Neither consideration makes low-level tracking insensitive to attention nor equates inattention with anesthesia. Awake attentional engagement was not measured, and attention was not independently manipulated across states, so its contribution to the observed differences cannot be isolated.

The clinical protocol and sampling design impose further limits. Anesthesia recordings were obtained during implantation surgery and wakefulness recordings later on the ward, confounding state with session order and recording context. The anesthetic regimen included adjunct medication and neuromuscular blockade, limiting attribution specifically to propofol. Clinical unresponsiveness defined the anesthesia condition, and auditory experience was not assessed trial by trial. Participant-level BIS and drug-exposure time series aligned to the analyzed epochs were unavailable for dose-response analyses. Electrode placement followed clinical requirements in patients with epilepsy, producing uneven regional coverage.

Future work could examine these joint response patterns across graded anesthetic states in a consistent recording setting, with time-aligned BIS and drug-exposure measurements. Behavioral probes at responsive levels of sedation and controlled attention tasks would help relate the neural measures to auditory performance and attentional engagement. Larger cohorts with broader anatomical coverage, where clinically feasible, and analyses that account for within-participant dependence would support assessment of the spatial patterns across individuals. Extending decoding to stimulus features beyond the acoustic envelope, comparing CEBRA with PCA, and varying embedding dimensionality would test which observed patterns generalize across stimulus descriptions and representation methods.

## Methods

### Participants

Thirteen adult patients with drug-resistant epilepsy (ages 20-49 years, 4 males, see Supplementary Table 1 for details) were recruited from the Shenzhen Second People’s Hospital, China. All participants were native Mandarin speakers and were right-handed according to the Edinburgh Handedness Inventory^39^. Patients underwent sEEG depth electrode implantation for presurgical evaluation to localize seizure foci. Electrode placement and all clinical procedures were determined solely by medical necessity and were not altered for research purposes. Data were acquired both during the implantation surgery under anesthesia and during subsequent monitoring on the medical ward. The study was approved by the Clinical Research Ethics Committee at Shenzhen Second People’s Hospital and the Institutional Review Board at New York University Shanghai (No. 2025-441-02PJ), and all participants provided written informed consent. The research was conducted in accordance with the Declaration of Helsinki, and participants received no compensation.

### Anesthesia protocol

General anesthesia was administered by an anesthesiologist as part of routine clinical care during sEEG electrode implantation. Standard intraoperative monitoring included bispectral index (BIS) and non-invasive arterial blood pressure (B20i; GE Healthcare, Helsinki, Finland). The anesthetic regimen for this dataset has been described previously^40^. Anesthesia was induced intravenously with etomidate (20 mg), sufentanil (20 µg), and rocuronium bromide (50 mg). Haloperidol (0.5 mg, i.v.) and dexamethasone (5 mg, i.v.) were also administered. Anesthesia was maintained using target-controlled infusions (TCI) of propofol and remifentanil. Propofol was delivered via an injection pump (CP-2200; SLGO Medical Technology, Beijing, China). Dexmedetomidine was continuously infused at 0.2 µg/kg/h. Anesthetic administration was adjusted to maintain stable deep anesthesia.

### Experiment design and stimuli

Participants were presented with an auditory oddball paradigm in both sessions of wakefulness and anesthesia (Fig. 1b). The stimuli comprised four different humming sounds, generated from a combination of two features: pitch contour (flat vs. rising tone) and intensity (60 dB SPL vs. 80 dB SPL). Each stimulus sequence consisted of five sounds. Within each sequence, four sounds were identical (“standard” stimulus) and one was acoustically distinct (“deviant” stimulus). The stimulus onset asynchrony (SOA) was fixed at 620 ms, resulting in a rate of approximately 1.61 Hz. The position of the deviant sound was pseudorandomized to occur in the second, third, or fourth position of the sequence. The experiment consisted of four blocks. In each block, one of the four sounds served as the standard, while the other three served as the possible deviants. This design ensured that each sound’s novelty was fully counterbalanced across the session. Each session contained 180 five-sound sequences (3 deviant types × 3 deviant positions × 5 repetitions × 4 blocks). The sequence order was pseudorandomized within each block. During wakefulness, participants were instructed to listen passively to the auditory sequences without making an overt response. Auditory stimuli were delivered binaurally via insert earphones (ER3C; Etymotic Research). Prior to each session, the volume was adjusted to a comfortable listening level as reported by the participants.

All core analyses pooled responses to standard and deviant sounds to characterize tracking at the sound-presentation rate. The data were originally collected for a study of mismatch and novelty responses in the same patients^40^. Here, we used their fixed sound-presentation rate to compare rhythmic auditory tracking during wakefulness and propofol-based anesthesia.

### sEEG data acquisition

The sEEG depth electrodes (Huake Hengsheng Medical, Beijing, China) featured a shaft diameter of 0.8 mm and contained 8–16 contacts with a center-to-center spacing of 3.5 mm. Across the patient cohort, a total of 151 electrodes (87 left hemisphere, 64 right hemisphere), comprising 2142 contacts (1230 left, 912 right), were implanted for clinical evaluation (Fig. 1a). The sEEG data were recorded with the 256-channel recording system (Nicolet Quantum; Natus Medical, San Carlos, CA, USA) at a sampling rate of 2048 Hz. Signals were online referenced to the average of two adjacent contacts located in deep white matter, chosen for their distance from gray matter and the suspected epileptogenic zone, in line with established methods^41^. Contacts were excluded from subsequent analysis if they were situated within the clinically determined epileptogenic zone, exhibited impedance exceeding 15 kΩ, or showed significant artifacts such as 50 Hz line noise.

### Electrode localization

We determined the anatomical localization of sEEG electrodes using a customized pipeline incorporating FreeSurfer^42^ and the Brainstorm toolbox^43^. The preoperative T1-weighted MRI scans (Prisma, 3T; Siemens, Munich, Germany) were co-registered with postoperative computed tomography (CT) scans (Somatom Definition Flash; Siemens, Munich, Germany) using an affine transformation implemented in SPM12. Following co-registration, we manually identified each electrode trajectory in the native CT volume by marking its tip and skull entry point. The three-dimensional coordinates of individual contacts along each electrode shaft were then automatically interpolated. Subsequently, we assigned each contact to an anatomical region using FreeSurfer’s automated cortical and subcortical parcellation scheme^44,45^. For group-level visualization, all patient-native electrode coordinates were normalized to the Montreal Neurological Institute (MNI) standard space (Fig. 1a).

### Signal preprocessing

Preprocessing used custom Python scripts and MNE-Python^46,47^. Channels identified by the clinical team as being located within the epileptic zone were excluded from all subsequent analyses. The raw data were first processed to remove electrical line noise and its harmonics. To enhance spatial specificity and mitigate the effects of volume conduction, the data were re-referenced using a bipolar montage (Fig. 1d). Following re-referencing, the data were band-pass filtered between 0.1 and 400 Hz (FIR filter). The preprocessed data were segmented into epochs, each comprising three consecutive five-sound sequences. Epochs extended from −0.2 to 9.3 s relative to the first sound onset in each group. Grouping the 180 stimulus sequences in this way yielded 60 analysis epochs per participant and recording state before artifact rejection. No baseline correction was applied; epochs were linearly detrended. Artifact rejection was performed at the epoch level. Any epoch containing activity exceeding an absolute amplitude threshold of 350 µV or with a variance greater than four standard deviations from the mean across all epochs for a given channel was discarded.

### Spectral power analysis

The power spectral density (PSD) for each epoch was computed using the Fast Fourier Transform (FFT) after applying a Hamming window to reduce spectral leakage. To accurately characterize oscillatory activity, we separated the periodic (peak) and aperiodic (1/f-like) components of the power spectrum. This was achieved by parameterizing each epoch’s PSD using the ‘fitting oscillations and one-over-f’(FOOOF) algorithm^48^. The power spectrum, P, for a given frequency, f, is modeled as the sum of an aperiodic component, L, and a series of Gaussian peaks, G, representing periodic oscillations:

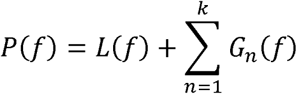

To assess the quality of spectral parameterization, we evaluated the goodness of fit for each channel by fitting the FOOOF model to the mean power spectrum across epochs and extracting the coefficient of determination (R²). No channels were excluded based on FOOOF fit quality (see Results and Supplementary Table 3). The aperiodic-adjusted power spectrum was obtained by subtracting the fitted aperiodic component from the original spectrum. To identify channels with a significant neural response at the stimulus frequency (1.61 Hz), we performed a one-tailed, paired t-test for each channel. This test compared the mean aperiodic-adjusted power at the target frequency bin against the average power in the two neighboring frequency bins^49^. The resulting p-values were corrected for multiple comparisons across all channels using the Bonferroni method, with a significance threshold of α = 0.05. This procedure was applied separately to wakefulness and anesthesia; a bipolar channel was classified as power responsive in a state only when it met this criterion in that state. To compare global auditory tracking power between wakefulness and anesthesia, we fitted a linear mixed-effects model^50^ (LME) with state (wakefulness vs. anesthesia) as a fixed effect and participant as a random intercept, accounting for the non-independence of multiple channels recorded from the same individual. The model was estimated using restricted maximum likelihood (REML).

### Phase-locking and temporal organization analysis

To assess temporal fidelity, the amplitude envelope of both the trial-averaged auditory stimulus and the trial-averaged sEEG signal for each channel was extracted using the Hilbert transform. These envelopes were then smoothed, down-sampled to 100 Hz, detrended, and band-pass filtered between 1.0 and 2.6 Hz to isolate the rhythm of the stimulus presentation. The Phase-Locking Value (PLV) was calculated as the magnitude of the mean vector of the phase differences between the two signals over time:

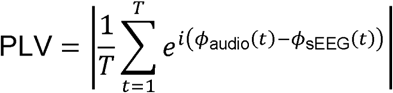

where T is the number of time points, and Φ_audio_(t) and Φ_sEEG_(t) are the instantaneous phases of the filtered stimulus and the sEEG amplitude envelopes, respectively. For each channel and state, onset phase concentration was calculated as:

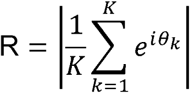

where θ_k_ is the instantaneous phase of the filtered sEEG amplitude envelope at the k^th^ included sound onset in the trial-averaged response, and K is the number of included onsets.

To test whether the timing of neural entrainment was altered by anesthesia, we compared the distributions of mean sEEG envelope phase angles at sound onsets between wakefulness and anesthesia^51^. This yielded two circular distributions of phase angles (wakefulness and anesthetized) across the population of responsive channels. The distribution of phase angles were compared using nonparametric Watson’s U^2^ test^52^.

### Spatial gradient and circular-linear correlation analysis

We investigated whether neural response magnitude (power) and timing (phase) followed specific spatial gradients across the brain. We defined two spatial metrics for each channel: Anterior-Posterior Axis: The MNI Y-coordinate; Surface distance from the auditory reference: We used the mesh-based shortest-path distance from an ipsilateral pmHG reference, motivated by the location of auditory core cortex^53^ and previous work on cortical spatial organization^54,55^. Reference coordinates were (−39, −22, 7) mm in the left hemisphere and (46, −24, 6) mm in the right hemisphere in the MNI space. The main power-distance analysis included 178 channels from 13 participants. All included channel locations, including noncortical sites, were projected to the nearest vertex of the ipsilateral fsaverage pial surface; for noncortical sites, this is a projected surface-distance descriptor rather than a path through the recorded tissue. Each reference point was projected to the nearest vertex on the fsaverage pial surface mesh. A weighted adjacency graph was constructed from the mesh triangulation, with edge weights equal to the Euclidean distance between connected vertices. Geodesic distances from the pmHG reference vertex to all other vertices were then computed using Dijkstra’s algorithm^56^, and each sEEG channel was assigned the geodesic distance of its nearest surface vertex.

Associations between stimulus-rate power and pmHG surface distance were quantified using standard linear correlations (Pearson’s r) between the spectral power (wakefulness and anesthesia) and both spatial metrics. To test whether the change in stimulus-rate power between states varied with distance from pmHG beyond participant-specific overall power shifts, we fitted an ordinary least-squares model to paired channel differences in power (anesthesia minus wakefulness), with participant-specific intercepts and distance expressed in units of 100 mm. The distance coefficient estimated the between-state slope contrast from within-participant spatial variation. Its 95% confidence interval was calculated using a participant-cluster bootstrap^57^ Statistical significance was assessed using a two-sided participant-level sign-flip test^58^.

To test whether the timing of neural entrainment followed a spatial gradient, we performed a circular-linear correlation analysis. We correlated the phase angle of each responsive sEEG channel with its corresponding normalized coordinate along the anterior-posterior axis (y-coordinate in MNI space). The correlation coefficient (ρ) was determined by^29^:

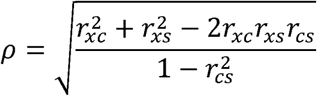

where r_xc_ is the correlation between the y-coordinate and cos cos(θ), r_xs_ is the correlation between the y-coordinate and sin(θ), and r_cs_ is the correlation between cos(θ) and sin(θ). The statistical significance was determined by converting it to a test statistic,nρ^2^, which follows a chi-squared distribution with two degrees of freedom, from which the p-value was derived (α=0.05). Additionally, to quantitatively model the progression of phase shifts across space, we fitted a circular-linear regression model for both wakefulness and anesthesia states, regressing the phase angle (θ) onto the spatial predictors (x).

### Region of interest (ROI) analysis

We analyzed channels’ power values/phase angles across eleven specific anatomical ROIs: Heschl’s gyrus (HG), Superior temporal gyrus (STG), Middle temporal gyrus (MTG), Inferior temporal gyrus (ITG), Fusiform gyrus (FG), Insula (INS), Rolandic operculum (RO), Inferior frontal gyrus (IFG), Postcentral gyrus (PoCG), Hippocampus (HIPP), and Putamen (PUT). To assess differences in power between states, we employed a stratified permutation test (n=10,000). For each ROI, the observed test statistic was the sum of Welch’s t-values^59^ across channels. A null distribution was generated by shuffling condition labels (wakefulness vs. anesthesia) within each channel independently. For phase angles, linear statistics were inappropriate due to the circular nature of the variable (0 to 2π). We therefore compared the phase distributions between wakefulness and anesthesia states using the Wheeler-Watson test^60^, a non-parametric circular statistic sensitive to differences in both mean angle and angular variance. All p values were corrected for multiple comparisons across the 11 ROIs using the Bonferroni correction.

### Visualization of electrode localization and neural responses

We visualized the spatial distribution of group-level sEEG channel coordinates (MNI space) and their corresponding power values/phase angles directly within a transparent “glass brain” volume, avoiding voxel-grid conversion or surface interpolation to preserve spatial precision. The visualization was generated using the “nilearn” library in Python^61^. In the response maps, each bipolar sEEG channel was represented by a node at the midpoint of its two contact coordinates in the MNI space. The power/phase magnitude was encoded by both the color (using a ‘hot’/’coolwarm’ colormap) and the size of the node (scaled proportionally to the value). These nodes were superimposed onto a standard MNI reference brain and displayed across lateral (left/right) and axial (top-down) views.

### Stimulus-aligned embedding modeling with CEBRA

We used CEBRA^24^ to learn three-dimensional neural embeddings conditioned on the acoustic envelope for decoding and consistency analyses (Fig. 4a). We configured CEBRA in a hypothesis-driven mode, conditioning the neural embedding on the continuous audio envelope. This approach incentivizes the model to project high-dimensional sEEG signals into a 3-dimensional latent space where neural states associated with similar acoustic features are clustered together, providing a compact representation in which to examine stimulus-related patterns of sEEG activity.

We employed a conditional CEBRA model configured with the ‘offset10-model’ architecture. The model was trained to map stimulus-rate power active channel sEEG data (both wakefulness and anesthesia state, pooled from all participants) to the 1-dimensional continuous audio envelope (behavioral variable). Before training, each channel’s data was normalized to the [0, 1] range, and the same scaling parameters were then applied to the validation and shuffled-control data. Key hyperparameters were fixed as follows: output dimension = 3; number of hidden units = 32; batch size = 512; learning rate = 1 × 10⁻□; temperature = 1; and time offsets = 10 samples. Training was performed for a maximum of 6000 iterations on an Nvidia RTX 4070 graphics card with 12 GB of memory.

### Acoustic-envelope decoding and model validation

We assessed model generalization using training-validation partitions of the epochs defined above. For each participant and recording state, the 60 epochs were pseudorandomly partitioned into 40 training epochs and 20 validation epochs. Model performance was evaluated using 30 repetitions of randomized three-fold partitioning, yielding 90 repeated model fits. We also trained separate “shuffled” control models. For each shuffled-control encoder, individual acoustic-envelope samples were randomly permuted across the combined two-state training timeline. These models used separately generated neural training and validation averages prepared with the same procedure as the original-label models. The downstream decoders were fitted and evaluated using the original time-aligned envelope labels. These controls assessed model performance after disrupting the original alignment between sEEG activity and acoustic-envelope labels.

Model performance was evaluated by training a k-nearest-neighbor (k-NN) decoder on the learned latent embeddings. We transformed the neural data into latent embeddings and trained the decoder to predict the corresponding acoustic-envelope values. The decoder was configured with “k=36” neighbors and used a cosine distance metric. For each validation set, the decoder was fit using the embeddings from the training set and the corresponding audio envelope labels. We then predicted the audio envelope from the held-out validation set embeddings. Decoding performance was assessed using two metrics: the coefficient of determination (R^2^) and the Median Absolute Error (MAE) between the predicted and true audio envelopes. These metrics were computed for both wakefulness and anesthesia conditions across all model fits, allowing for statistical comparison against the chance-level baseline derived from the shuffled control models. Finally, three-dimensional embeddings were generated by transforming the neural data through the trained CEBRA models, illustrating the learned stimulus-aligned representations.

To test whether validation performance depended on the amount of available training data, we performed a data titration analysis. Goodness of fit (GoF) was computed separately for each state using the neural trajectory and corresponding envelope labels. It estimates InfoNCE loss and converts it to GoF in bits. Models were trained using 20%, 40%, 60%, 80%, and 100% of the training epochs within each cross-validation fold (10 random seeds × 3 folds per fraction; 150 additional models total). Validation performance (R² and GoF) was evaluated on the same held-out validation set at each data fraction. The generalization gap (training minus validation metric) was computed for each model to characterize how the train-validation discrepancy scaled with data amount.

### Embedding consistency and channel-embedding associations

We assessed embedding consistency across repeated model fits using the “consistency_score” metric provided by CEBRA. Each fitted encoder was applied to its validation responses, and consistency was evaluated separately for wakefulness and anesthesia and for their respective shuffled controls. The metric quantifies correspondence between embeddings using linear-regression R² at corresponding time points. The regression is fitted and scored on the same paired embedding rows. Higher scores indicate greater reproducibility of the learned embeddings under the tested conditions. Differences between conditions were assessed using Mann-Whitney U tests^62^.

We quantified marginal associations between individual channels and the CEBRA embeddings. A separate linear regression related each channel’s activity to the three-dimensional embedding coordinates. Each regression was fitted and scored on the same encoder-validation trajectory, yielding a descriptive channel-embedding R². These channel-wise R² values quantify associations with the embedding coordinates; they may include variance shared with other channels and do not identify unique or causal contributions. This procedure was performed separately for wakefulness and anesthesia. The R² values were mapped to the anatomical locations of the channels.

### Simulation analyses

We used two simulations to characterize how controlled amplitude, phase, and spatial rescaling manipulations affected the measured responses.

Simulation 1: Effects of amplitude and phase manipulations. We generated synthetic ERP trials (60 trials, 9300 samples at 1000 Hz) under four scenarios that independently manipulated response amplitude (high: 2.0; low: 0.05) and phase alignment (locked: jitter = 0.2 rad; spread: phases evenly distributed over 0–2π). Each trial contained a periodic Gaussian ERP train at 1.61 Hz embedded in 1/f pink noise (amplitude = 0.15). Spectral power was measured using FOOOF on the mean PSD across trials, extracting the aperiodic-corrected power at 1.61 Hz. PLV was computed following the same method used in the paper. Fifty independent repetitions per scenario were generated, and cross-effects were quantified as the percentage of the main effect explained by the orthogonal manipulation. Statistics were computed using two-sample t-tests.

Simulation 2: Spatial correlations under response rescaling. To characterize the effects of response rescaling on distance-response correlations, we generated 200 synthetic channels with geodesic-like distances calibrated to match the observed wakefulness correlation. We applied uniform positive scaling (factors 0.1-1.0) and distance-dependent attenuation separately. Under uniform positive scaling, the correlation is unchanged: r(X, aY) = r(X, Y) for a > 0.

## Acknowledgments

We thank Xinjing Li, Wenqiang Zhang, Xiao Li, Lanhui Huang, Minhua Zhang, and Wenjing Wang for their assistance on data collection. We thank Shaoqi Pan and Yixiang Liu for their constructive advice on data analysis and visualization.

## Data availability

Derived data supporting selected summary-statistical analyses in this study are available on Zenodo at https://doi.org/10.5281/zenodo.23655785. Raw sEEG recordings are available from the corresponding author upon request.

## Code availability

Python scripts for reproducing the deposited summary-statistical analyses are available on Zenodo at https://doi.org/10.5281/zenodo.23655785. Additional analysis code is available from the corresponding author upon request.

## Author contributions

H.Z., X.Tian, and X.T. conceptualized the study. H.Z. and C.Y. performed experiments. H.Z., C.Y., W.Z., Y.W., S.L., and Z.H. collected the data. H.Z. analyzed the data. H.Z. and X.T. wrote the manuscript. All authors reviewed the manuscript.

## References

1. Alkire, M. T., Hudetz, A. G. & Tononi, G. Consciousness and anesthesia. Science 322, 876–880 (2008).

2. Brown, E. N., Lydic, R. & Schiff, N. D. General anesthesia, sleep, and coma. N. Engl. J. Med. 363, 2638–2650 (2010).

3. Hudetz, A. G. General anesthesia and human brain connectivity. Brain Connect. 2, 291–302 (2012).

4. Davis, M. H. et al. Dissociating speech perception and comprehension at reduced levels of awareness. Proc. Natl. Acad. Sci. 104, 16032–16037 (2007).

5. Plourde, G. et al. Cortical processing of complex auditory stimuli during alterations of consciousness with the general anesthetic propofol. Anesthesiology 104, 448–457 (2006).

6. Uhrig, L., Janssen, D., Dehaene, S. & Jarraya, B. Cerebral responses to local and global auditory novelty under general anesthesia. Neuroimage 141, 326–340 (2016).

7. Bonhomme, V., Plourde, G., Meuret, P., Fiset, P. & Backman, S. B. Auditory steady-state response and bispectral index for assessing level of consciousness during propofol sedation and hypnosis. Anesth. Analg. 91, 1398–1403 (2000).

8. Plourde, G. & Picton, T. W. Human auditory steady-state response during general anesthesia. Anesth. Analg. 71, 460–468 (1990).

9. Krom, A. J. et al. Anesthesia-induced loss of consciousness disrupts auditory responses beyond primary cortex. Proc. Natl. Acad. Sci. 117, 11770–11780 (2020).

10. Nourski, K. V. et al. Auditory Predictive Coding across Awareness States under Anesthesia: An Intracranial Electrophysiology Study. J. Neurosci. 38, 8441–8452 (2018).

11. Nourski, K. V. et al. Cortical responses to vowel sequences in awake and anesthetized states: a human intracranial electrophysiology study. Cereb. Cortex 31, 5435–5448 (2021).

12. Näätänen, R., Kujala, T. & Winkler, I. Auditory processing that leads to conscious perception: a unique window to central auditory processing opened by the mismatch negativity and related responses. Psychophysiology 48, 4–22 (2011).

13. Näätänen, R. The role of attention in auditory information processing as revealed by event-related potentials and other brain measures of cognitive function. Behav. Brain Sci. 13, 201–233 (1990).

14. Ding, N. & Simon, J. Z. Neural coding of continuous speech in auditory cortex during monaural and dichotic listening. J. Neurophysiol. 107, 78–89 (2012).

15. Ding, N. et al. Attention is required for knowledge-based sequential grouping: insights from the integration of syllables into words. J. Neurosci. 38, 1178–1188 (2018).

16. Batterink, L. J. & Paller, K. A. Statistical learning of speech regularities can occur outside the focus of attention. Cortex 115, 56–71 (2019).

17. Hayat, H. et al. Reduced neural feedback signaling despite robust neuron and gamma auditory responses during human sleep. Nat. Neurosci. 25, 935–943 (2022).

18. Alnes, S. L. et al. Auditory responses in the temporal lobe are modulated by slow waves of sleep. Prog. Neurobiol. 255, 102843 (2025).

19. Henry, M. J., Herrmann, B. & Obleser, J. Entrained neural oscillations in multiple frequency bands comodulate behavior. Proc. Natl. Acad. Sci. 111, 14935–14940 (2014).

20. Teng, X., Tian, X., Rowland, J. & Poeppel, D. Concurrent temporal channels for auditory processing: Oscillatory neural entrainment reveals segregation of function at different scales. PLoS Biol. 15, e2000812 (2017).

21. Lakatos, P., Gross, J. & Thut, G. A new unifying account of the roles of neuronal entrainment. Curr. Biol. 29, R890–R905 (2019).

22. Obleser, J. & Kayser, C. Neural entrainment and attentional selection in the listening brain. Trends Cogn. Sci. 23, 913–926 (2019).

23. Ding, N. & Simon, J. Z. Cortical entrainment to continuous speech: functional roles and interpretations. Front. Hum. Neurosci. 8, 311 (2014).

24. Schneider, S., Lee, J. H. & Mathis, M. W. Learnable latent embeddings for joint behavioural and neural analysis. Nature 617, 360–368 (2023).

25. Kösem, A., Gramfort, A. & Van Wassenhove, V. Encoding of event timing in the phase of neural oscillations. Neuroimage 92, 274–284 (2014).

26. Zoefel, B. & VanRullen, R. EEG oscillations entrain their phase to high-level features of speech sound. Neuroimage 124, 16–23 (2016).

27. Fisher, N. I., Lewis, T. & Embleton, B. J. Statistical Analysis of Spherical Data. (Cambridge university press, 1993).

28. Van Diepen, R. M. & Mazaheri, A. The caveats of observing inter-trial phase-coherence in cognitive neuroscience. Sci. Rep. 8, 2990 (2018).

29. Zar, J. H. Biostatistical Analysis. (Pearson Education Limited, Harlow, 2014).

30. Gallego, J. A., Perich, M. G., Miller, L. E. & Solla, S. A. Neural manifolds for the control of movement. Neuron 94, 978–984 (2017).

31. Stringer, C., Pachitariu, M., Steinmetz, N., Carandini, M. & Harris, K. D. High-dimensional geometry of population responses in visual cortex. Nature 571, 361–365 (2019).

32. Vyas, S., Golub, M. D., Sussillo, D. & Shenoy, K. V. Computation through neural population dynamics. Annu. Rev. Neurosci. 43, 249–275 (2020).

33. Chung, S. & Abbott, L. F. Neural population geometry: An approach for understanding biological and artificial neural networks. Curr. Opin. Neurobiol. 70, 137–144 (2021).

34. Banks, M. I. et al. Functional geometry of auditory cortical resting state networks derived from intracranial electrophysiology. PLoS Biol. 21, e3002239 (2023).

35. Zhang, Y., Leonard, M. K., Bhaya-Grossman, I., Gwilliams, L. & Chang, E. F. Human cortical dynamics of auditory word form encoding. Neuron 114, 167–180 (2026).

36. Eisen, A. J. et al. Propofol anesthesia destabilizes neural dynamics across cortex. Neuron 112, 2799–2813 (2024).

37. Ensel, S. et al. Transient brain activity dynamics discriminate levels of consciousness during anesthesia. Commun. Biol. 7, 716 (2024).

38. Jang, H., Mashour, G. A., Hudetz, A. G. & Huang, Z. Measuring the dynamic balance of integration and segregation underlying consciousness, anesthesia, and sleep in humans. Nat. Commun. 15, 9164 (2024).

39. Oldfield, R. C. The assessment and analysis of handedness: the Edinburgh inventory. Neuropsychologia 9, 97–113 (1971).

40. Han, Z. et al. Automatic binding of basic sensory features requires consciousness. bioRxiv 10.64898/2025.12.10.693567 (2025) doi:10.64898/2025.12.10.693567.

41. Xiao, L. et al. Automatic localization of seizure onset zone from high-frequency SEEG signals: A preliminary study. IEEE J. Transl. Eng. Health Med. 9, 1–10 (2021).

42. Fischl, B. FreeSurfer. Neuroimage 62, 774–781 (2012).

43. Tadel, F., Baillet, S., Mosher, J. C., Pantazis, D. & Leahy, R. M. Brainstorm: A user-friendly application for MEG/EEG analysis. Comput. Intell. Neurosci. 2011, 879716 (2011).

44. Destrieux, C., Fischl, B., Dale, A. & Halgren, E. Automatic parcellation of human cortical gyri and sulci using standard anatomical nomenclature. Neuroimage 53, 1–15 (2010).

45. Rolls, E. T., Huang, C.-C., Lin, C.-P., Feng, J. & Joliot, M. Automated anatomical labelling atlas 3. Neuroimage 206, 116189 (2020).

46. Gramfort, A. et al. MEG and EEG data analysis with MNE-Python. Front. Neurosci. 7, 267 (2013).

47. Gramfort, A. et al. MNE software for processing MEG and EEG data. neuroimage 86, 446–460 (2014).

48. Donoghue, T. et al. Parameterizing neural power spectra into periodic and aperiodic components. Nat. Neurosci. 23, 1655–1665 (2020).

49. Ding, N., Melloni, L., Zhang, H., Tian, X. & Poeppel, D. Cortical tracking of hierarchical linguistic structures in connected speech. Nat. Neurosci. 19, 158–164 (2016).

50. Baayen, R. H., Davidson, D. J. & Bates, D. M. Mixed-effects modeling with crossed random effects for subjects and items. J. Mem. Lang. 59, 390–412 (2008).

51. Repp, B. H. Sensorimotor synchronization: A review of the tapping literature. Psychon. Bull. Rev. 12, 969–992 (2005).

52. Watson, G. S. Goodness-of-fit tests on a circle. II. Biometrika 49, 57–63 (1962).

53. Norman-Haignere, S. V. & McDermott, J. H. Neural responses to natural and model-matched stimuli reveal distinct computations in primary and nonprimary auditory cortex. PLoS Biol. 16, e2005127 (2018).

54. Margulies, D. S. et al. Situating the default-mode network along a principal gradient of macroscale cortical organization. Proc. Natl. Acad. Sci. 113, 12574–12579 (2016).

55. Royer, J. et al. Human cortical dynamics reflect graded contributions of local geometry and network topography. Nat. Commun. 17, 1002 (2026).

56. Dijkstra, E. W. A note on two problems in connexion with graphs. in Edsger Wybe Dijkstra: his life, work, and legacy 287–290 (2022).

57. Field, C. A. & Welsh, A. H. Bootstrapping clustered data. J. R. Stat. Soc. Ser. B Stat. Methodol. 69, 369–390 (2007).

58. Winkler, A. M., Ridgway, G. R., Webster, M. A., Smith, S. M. & Nichols, T. E. Permutation inference for the general linear model. Neuroimage 92, 381–397 (2014).

59. Welch, B. L. The significance of the difference between two means when the population variances are unequal. Biometrika 29, 350–362 (1938).

60. Wheeler, S. & Watson, G. S. A distribution-free two-sample test on a circle. Biometrika 51, 256–257 (1964).

61. Abraham, A. et al. Machine learning for neuroimaging with scikit-learn. Front. Neuroinformatics 8, 14 (2014).

62. Mann, H. B. & Whitney, D. R. On a test of whether one of two random variables is stochastically larger than the other. Ann. Math. Stat. 18, 50–60 (1947).

