## supplementary_information for "Auditory tracking magnitude, temporal alignment, and anatomical distribution during wakefulness and propofol-based anesthesia"

#### Supplementary Fig. 1

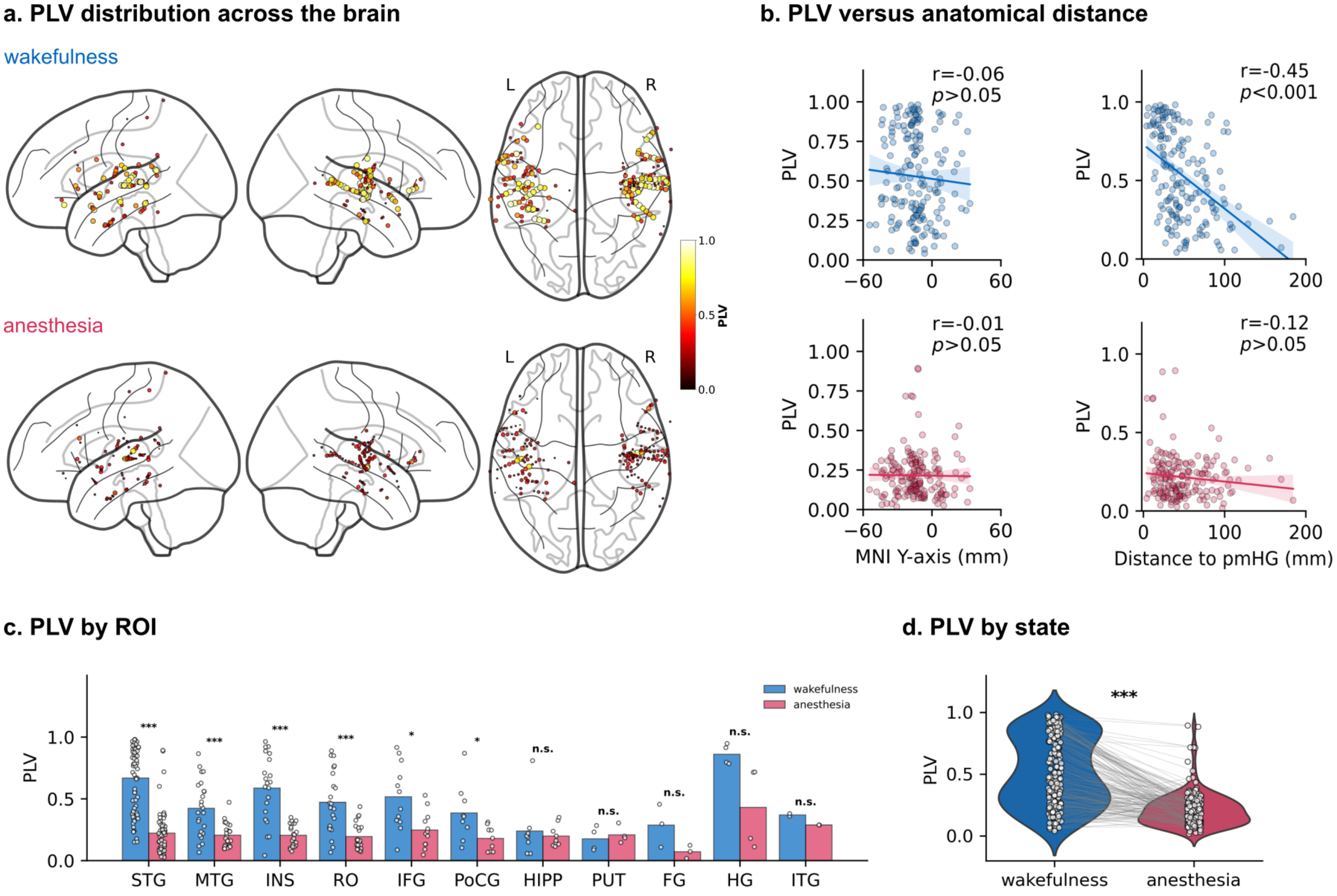


**Supplementary Fig. 1. Phase-locking value (PLV) and its spatial distribution during wakefulness and anesthesia.** (a) Spatial distribution of PLV across the analyzed channels, projected onto a glass brain in MNI space. Node color and size indicate PLV during wakefulness (top) and anesthesia (bottom). (b) Associations between PLV and anterior-posterior position (MNI Y-coordinate; left) or cortical surface distance from the pmHG reference (right). Solid lines show linear regression fits, and shading shows 95% confidence intervals. PLV was negatively associated with distance from pmHG during wakefulness ($r = -0.45$, $p < 0.001$); the association during anesthesia was weaker and not statistically significant ($r = -0.12$, $p > 0.05$). Associations with MNI Y-coordinate were not statistically significant in either state. (c) Mean PLV by anatomical region, with individual channel values overlaid. The plotted paired comparisons indicate lower PLV during anesthesia in STG, MTG, INS, RO, IFG, and PoCG. The HG comparison is marked not significant, as are HIPP, PUT, FG, and ITG. Significance annotations correspond to two-sided paired Wilcoxon signed rank tests: ***: $p < 0.001$; **: $p < 0.01$; *: $p < 0.05$; n.s., not significant. (d) PLV distributions across the analyzed channels in each state, with lines connecting paired channel measurements. Overall envelope PLV was lower during anesthesia than during wakefulness ($p < 0.001$).

### Supplementary Fig. 2

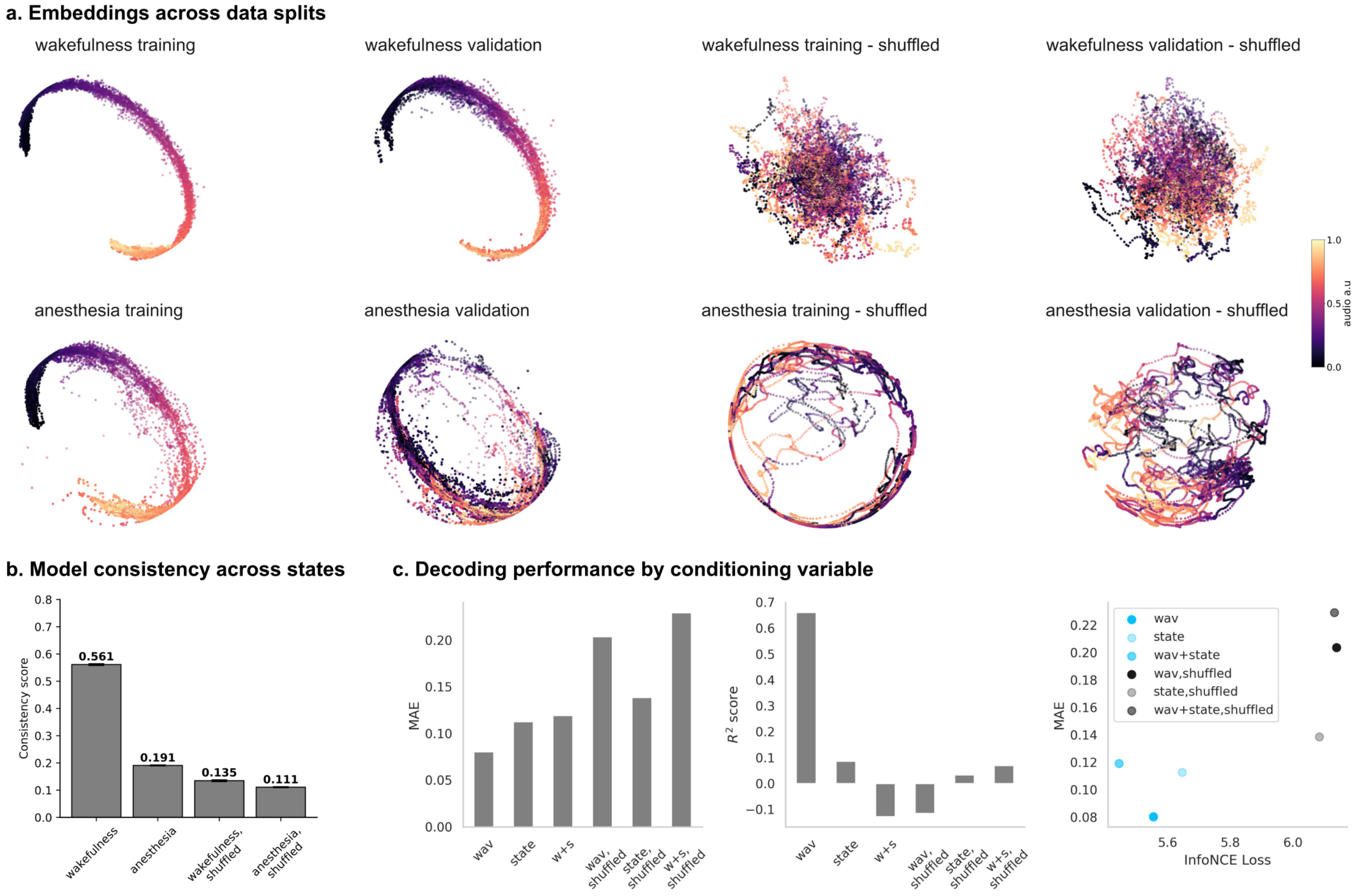


**Supplementary Fig. 2. Validation of CEBRA embeddings.** (a) Representative three-dimensional embeddings for training and validation data during wakefulness (top row) and anesthesia (bottom row), with their respective shuffled controls in the right columns. Colors indicate the acoustic envelope. (b) Embedding consistency across repeated model fits (mean ± s.e.m.; 90 fits). Consistency was evaluated separately for each state and its shuffled control using linear-regression $R^2$ at corresponding time points in the validation embeddings, with the regression fitted and scored on the same paired rows. Mean consistency scores were 0.561 for wakefulness, 0.191 for anesthesia, 0.135 for wakefulness shuffled, and 0.111 for anesthesia shuffled. (c) Decoding performance by conditioning variable. Left and middle: median absolute error (MAE) and decoding $R^2$ for models conditioned on the continuous acoustic envelope (wav), state labels (state), or both (w+s), alongside shuffled-label controls. Right: relationship between InfoNCE loss and MAE. Blue shades identify the original-label models, and gray/black symbols identify shuffled controls. Acoustic-envelope conditioning gave the lowest decoding error and highest decoding $R^2$ among the configurations shown. For the acoustic-envelope controls, encoders were trained with permuted envelope labels; downstream decoders were fitted and evaluated using the original time-aligned labels (Methods).

#### Supplementary Fig. 3

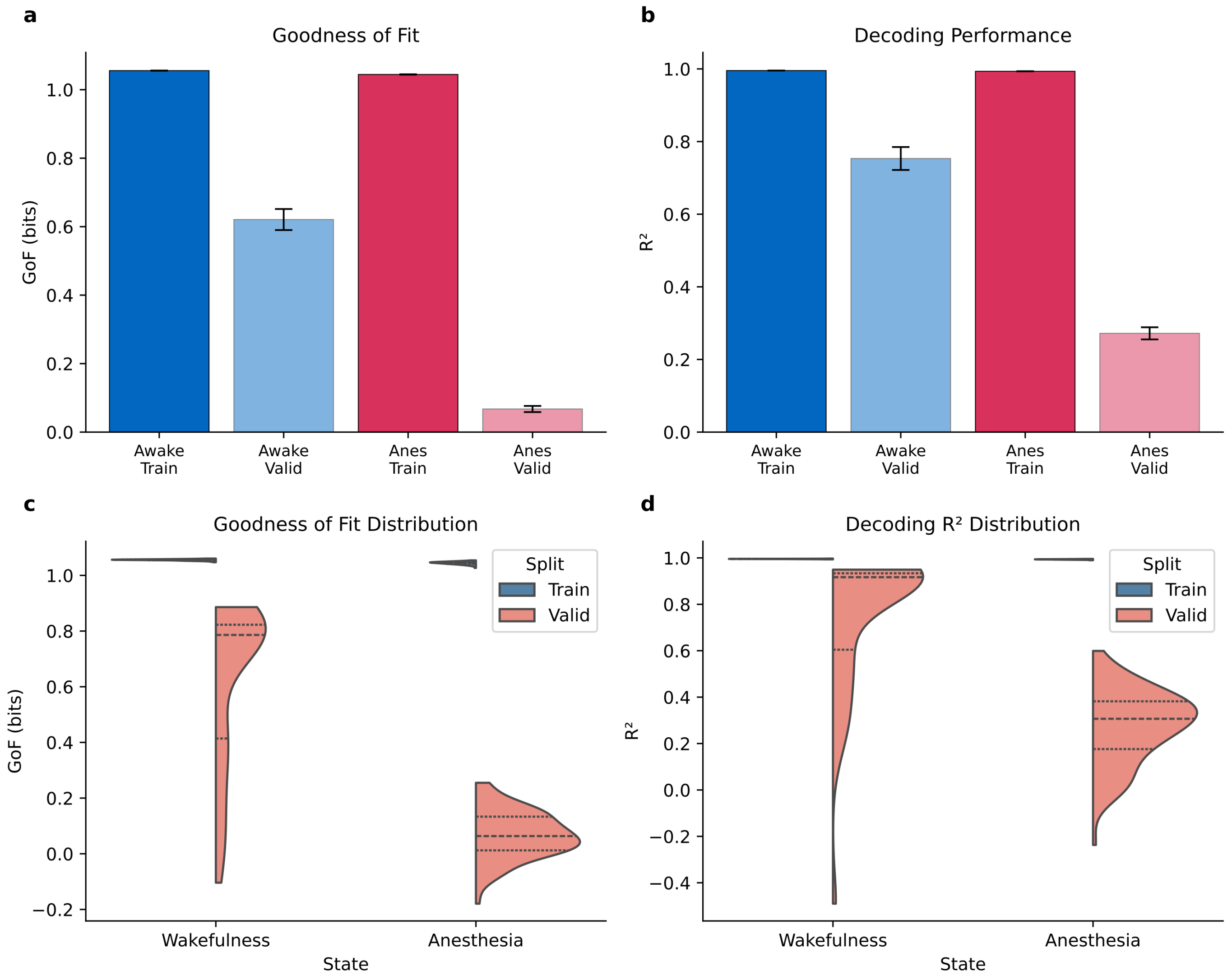


**Supplementary Fig. 3. Network training performance and generalization gap for wakefulness and anesthesia.** (a-b) Mean goodness of fit (GoF, in bits; a) and acoustic-envelope decoding $R^2$ (b) for training and validation splits (mean ± s.e.m.; 90 model fits from 30 repetitions of three-fold partitioning). Both states showed high training performance, with lower validation performance during anesthesia. (c-d) Distributions of GoF (c) and decoding $R^2$ (d) across the 90 model fits. The training-minus-validation GoF gap was 0.435 ± 0.031 bits during wakefulness and 0.977 ± 0.009 bits during anesthesia (mean ± s.e.m.). Corresponding $R^2$ gaps were 0.242 ± 0.032 and 0.722 ± 0.017. Paired Wilcoxon signed-rank comparisons showed a larger generalization gap during anesthesia ($p < 0.001$).

#### Supplementary Fig. 4

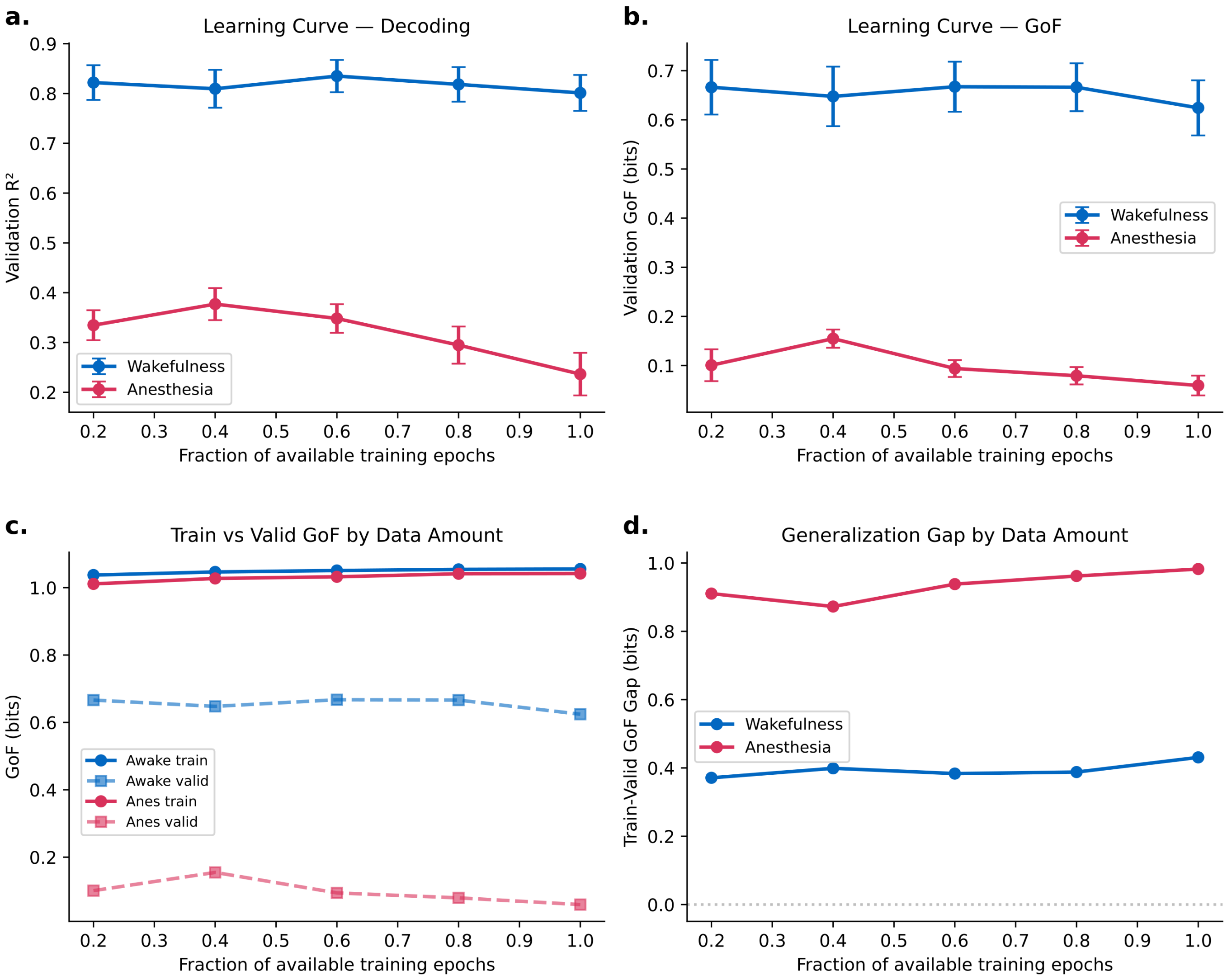


**Supplementary Fig. 4. Learning curve analysis: validation performance as a function of training data amount.** (a-b) Validation $R^2$ (a) and validation GoF in bits (b) plotted against the fraction of available training epochs used (20%, 40%, 60%, 80%, and 100%). Wakefulness validation performance remained within a similar range across fractions. Anesthesia validation performance remained lower at every fraction and did not improve consistently as more training data were used. (c) Training and validation GoF, showing persistently high training performance in both states and lower validation performance during anesthesia. (d) Generalization gap (training minus validation GoF). The gap was larger during anesthesia across all training fractions; after decreasing from 20% to 40%, it increased from 40% to 100%. The wakefulness gap remained smaller across all fractions. Models were trained with 10 random seeds × 3 cross-validation folds per fraction (150 additional model fits total), with the same held-out validation set within each fold at every fraction. Error bars in a and b show ± s.e.m.

#### Supplementary Fig. 5

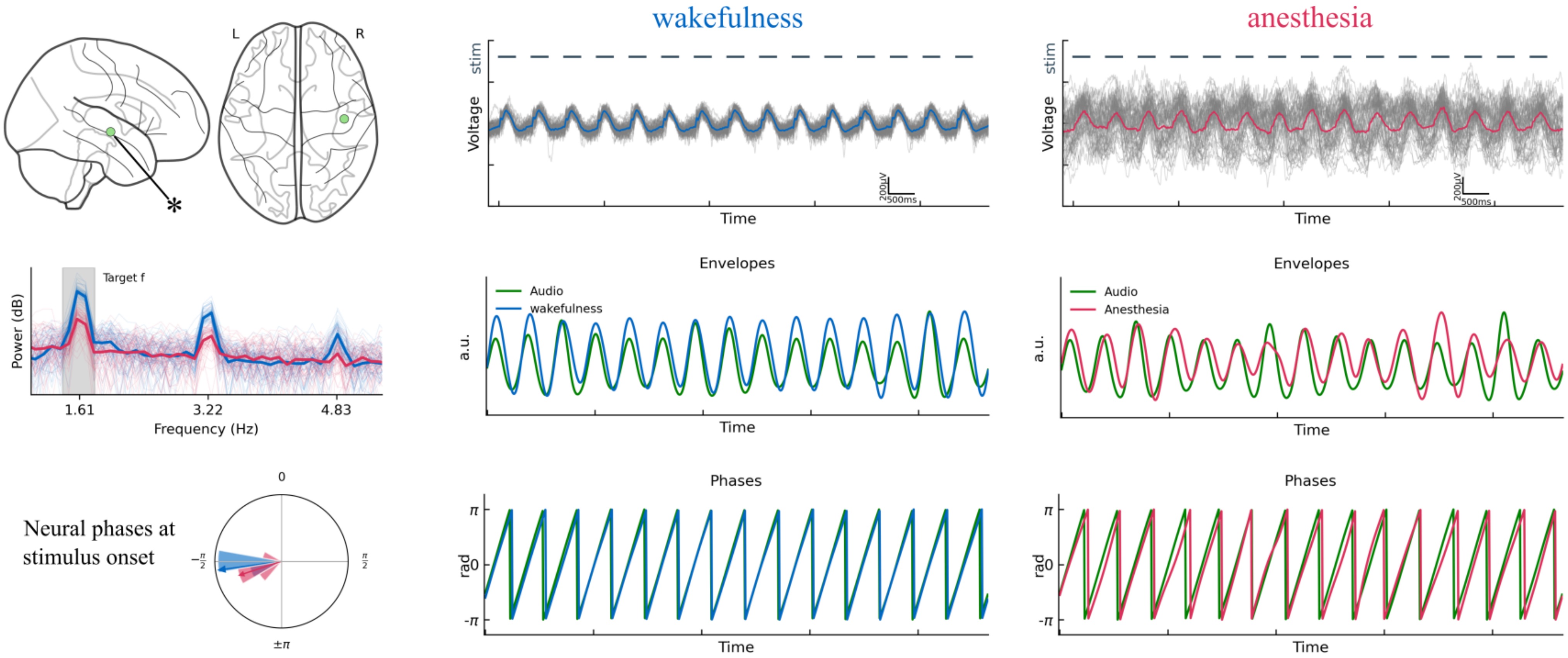


**Supplementary Fig. 5. Representative illustration of auditory tracking during wakefulness and anesthesia.** Data are shown from a representative sEEG channel located in the right superior temporal gyrus (green dot; top-left panels show lateral and axial glass brain views). Left column, middle: Power spectra for all trials (thin lines) and the mean (bold) during wakefulness (blue) and anesthesia (red). A prominent peak at the stimulus frequency (1.61 Hz, shaded) is visible during wakefulness, with attenuated power under anesthesia. Left column, bottom: Polar histograms of neural phase angles at stimulus onset, showing a concentrated distribution during wakefulness (blue) and anesthesia (red). Middle and right columns, top: Raw voltage traces across trials (gray) and the trial-averaged response (colored) during wakefulness (blue) and anesthesia (red). Dashed lines indicate the stimulus sequence. Middle and right columns, middle: Amplitude envelopes of the audio stimulus (green) and the neural signal (blue/red), band-pass filtered at the stimulus rate. Middle and right columns, bottom: Instantaneous phase time series of the audio (green) and neural (blue/red) signals.

### Supplementary Fig. 6

#### a. Amplitude and phase manipulations

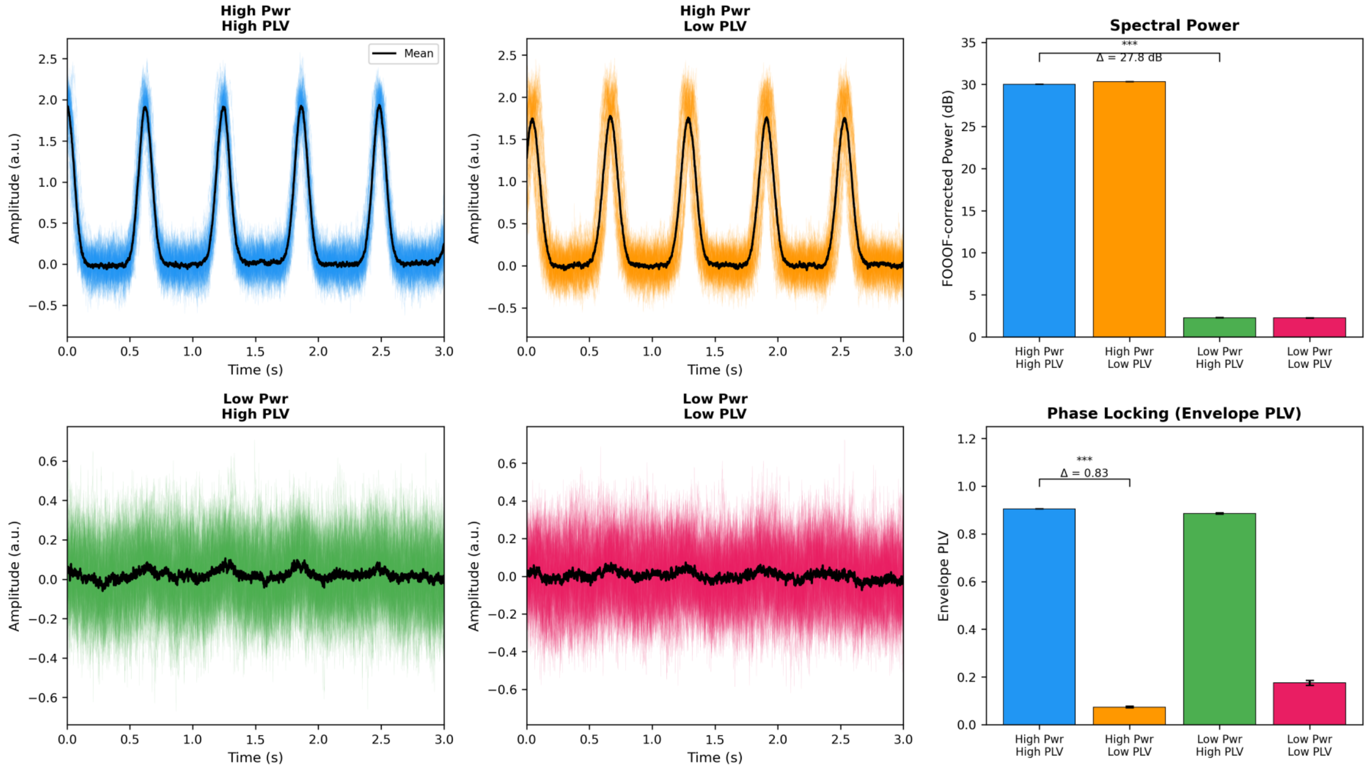


#### b. Power spectral density overlays

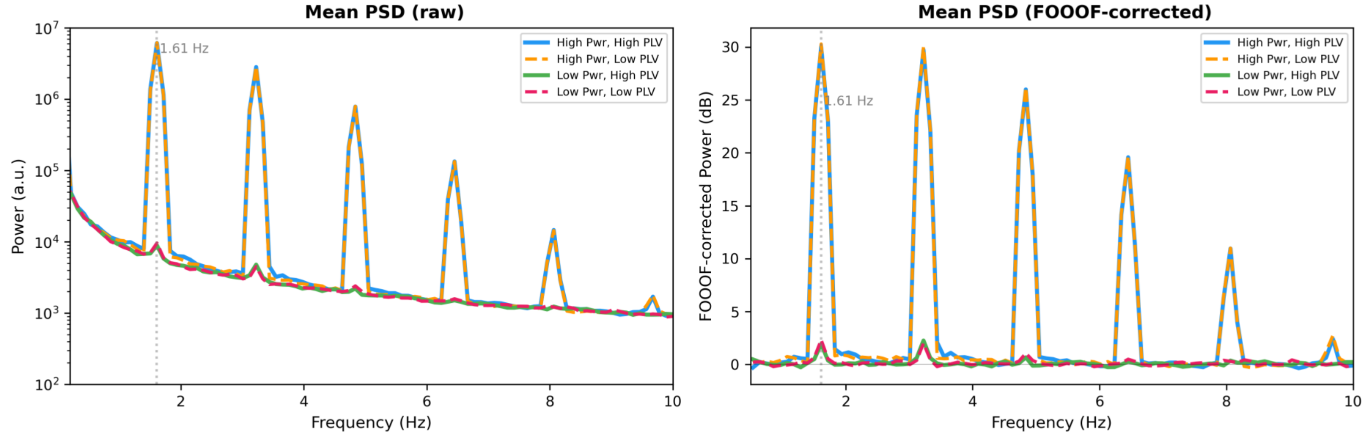


#### c. Spatial gradient and uniform scaling

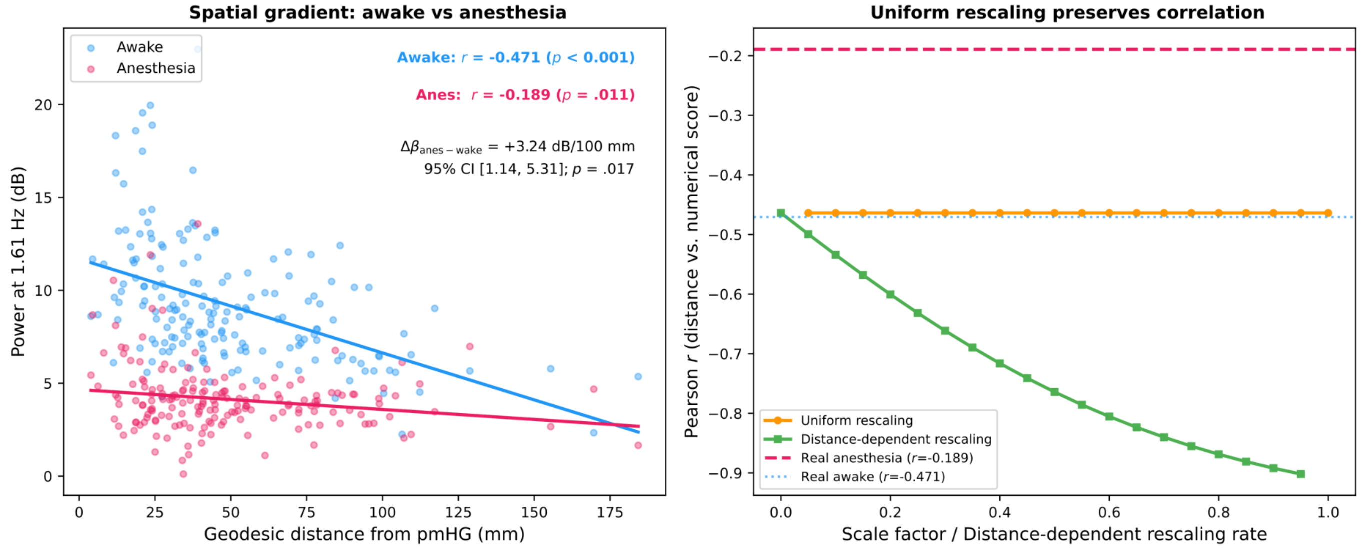


Supplementary Fig. 6 legend follows on the next page.

#### Supplementary Fig. 6 legend

**Supplementary Fig. 6. Simulated effects of response amplitude and phase alignment on power, phase measures, and distance-response relationships.** (a) Effects of amplitude and phase manipulations. Synthetic trials were generated under four combinations of high or low response amplitude and high or low phase alignment. The four panels on the left show example simulated signal traces over time; colored traces represent individual trials, and black traces show the trial averages. The panels on the right summarize FOOOF-corrected power at the stimulus rate (top) and envelope PLV (bottom) across 50 independent repetitions per scenario. The amplitude manipulation changed power by 27.75 dB and PLV by 0.020; the PLV change was 2.4% of the main effect of the phase manipulation. The phase manipulation changed PLV by 0.831 and power by 0.32 dB; the power change was 1.1% of the main effect of the amplitude manipulation. These results describe the relative sensitivities of the two measures under the tested signal model and parameter settings. (b) Power spectral density overlay for the four simulation scenarios: raw mean PSD on the left and FOOOF-corrected mean PSD on the right. The stimulus frequency is 1.61 Hz. Higher-amplitude conditions show a larger stimulus-rate peak under both phase-alignment scenarios. (c) Spatial correlations under response rescaling. Left: observed spectral power versus surface distance from the pmHG reference during wakefulness ($r = -0.471$, $p < 0.001$) and anesthesia ($r = -0.189$, $p = 0.011$). The participant-fixed-intercept slope contrast, estimated from paired channel power differences in 13 participants, was +3.24 dB/100 mm (anesthesia minus wakefulness; 95% participant-cluster bootstrap confidence interval, 1.14 to 5.31; two-sided participant-level sign-flip $p = 0.017$). Right: uniform positive scaling by factors from 0.1 to 1.0 preserved the correlation, whereas the tested distance-dependent attenuation progressively strengthened the negative correlation.

#### Supplementary Table 1

| Patient Code | Sex | Age (y) | Hand | Seizure Zone | Contacts (Rec./Imp.) | Electrodes (Rec./Imp.) |
|---|---|---|---|---|---|---|
| sz02 | M | 22 | RH | DCG.R | 177/212 | 13/13 |
| sz03 | M | 23 | RH | PrCG.L, SMA.L, PoCG.L | 127/212 | 14/15 |
| sz04 | F | 32 | RH | PHG.L, HIPP.L | 184/213 | 15/15 |
| sz05 | F | 34 | RH | MTG.R, INS.R, HIPP.R | 69/154 | 11/11 |
| sz07 | M | 32 | RH | PHG.R, HIPP.R | 126/164 | 11/11 |
| sz08 | F | 38 | RH | MTG.L&R, STG.R, INS.R | 69/159 | 10/12 |
| sz09 | F | 49 | RH | INS.L, STG.L | 107/151 | 12/12 |
| sz10 | M | 20 | RH | IPG.L, SMG.L | 130/187 | 11/13 |
| sz11 | F | 36 | RH | ACC.L, SFG.L | 153/175 | 12/12 |
| sz13 | F | 34 | RH | PCC.L | 74/119 | 7/8 |
| sz15 | F | 29 | RH | INS.L, RO.L | 82/103 | 7/7 |
| sz16 | F | 20 | RH | PHG.L, INS.L, HIPP.L | 122/163 | 12/12 |
| sz17 | F | 25 | RH | ITG.R, MTG.R | 105/130 | 10/10 |

**Supplementary Table 1. Participant demographics and electrode coverage.** Demographic and clinical characteristics of the 13 participants included in the study. The table details patient code, sex, age, handedness, and the clinically identified seizure onset zone. The "Contacts" and "Electrodes" columns quantify the coverage, displaying the ratio of channels analyzed (after artifact rejection and exclusion of the epileptic zone) relative to the total number implanted. Abbreviations: F, female; M, male; RH, right-handed; L, left hemisphere; R, right hemisphere; Rec., Recorded; Imp., Implanted; ACC, anterior cingulate cortex; DCG, middle cingulate & paracingulate gyri; HG, Heschl's gyrus; HIPP, hippocampus; INS, insula; IPG, inferior parietal gyrus; ITG, inferior temporal gyrus; MFG, middle frontal gyrus; MTG, middle temporal gyrus; PCC, posterior cingulate cortex; PHG, parahippocampal gyrus; PoCG, postcentral gyrus; PrCG, precentral gyrus; RO, Rolandic operculum; SFG, superior frontal gyrus; SMA, supplementary motor area; SMG, supramarginal gyrus; STG, superior temporal gyrus.

#### Supplementary Table 2

##### Per-ROI overlap

| ROI | wakefulness (wake) | anesthesia (anes) | both | only wake | only anes | anes in wake (%) | anes ⊆ wake |
|---|---|---|---|---|---|---|---|
| STG | 63 | 10 | 10 | 53 | 0 | 100 | true |
| MTG | 24 | 1 | 1 | 23 | 0 | 100 | true |
| INS | 21 | 2 | 2 | 19 | 0 | 100 | true |
| RO | 21 | 2 | 1 | 20 | 1 | 50 | no |
| IFG | 11 | 1 | 0 | 11 | 1 | 0 | no |
| PoCG | 9 | 0 | 0 | 9 | 0 | - | - |
| HIPP | 7 | 2 | 0 | 7 | 2 | 0 | no |
| PUT | 4 | 0 | 0 | 4 | 0 | - | - |
| FG | 3 | 0 | 0 | 3 | 0 | - | - |
| HG | 3 | 4 | 3 | 0 | 1 | 75 | no |
| ITG | 2 | 0 | 0 | 2 | 0 | - | - |

**Supplementary Table 2. Distribution and overlap of power-responsive bipolar sEEG channels.** Bipolar channels were classified by comparing aperiodic-adjusted power at 1.61 Hz with the mean power in the two neighboring frequency bins using a one-tailed paired t-test, with Bonferroni correction across all channels and a significance threshold of $\alpha = 0.05$, applied separately in each state (Methods). For each region, the table reports the number of channels responsive during wakefulness, anesthesia, both states, only wakefulness, and only anesthesia. “Anes in wake (%)” is the percentage of anesthesia-responsive channels also responsive during wakefulness. The final column indicates whether the anesthesia-responsive channels are a subset of the wakefulness-responsive channels in that region. Abbreviations: FG, fusiform gyrus; HG, Heschl’s gyrus; HIPP, hippocampus; IFG, inferior frontal gyrus; INS, insula; ITG, inferior temporal gyrus; MTG, middle temporal gyrus; PoCG, postcentral gyrus; PUT, putamen; RO, Rolandic operculum; STG, superior temporal gyrus.

#### Supplementary Table 3

| Participant | Channel N | Wakefulness $R^2$ (mean ± SD) | Wakefulness $R^2 < 0.9$ N_ch | Anesthesia $R^2$ (mean ± SD) | Anesthesia $R^2 < 0.9$ N_ch |
|---|---|---|---|---|---|
| sz02 | 177 | 0.978 ± 0.019 | 1 | 0.988 ± 0.003 | 0 |
| sz03 | 127 | 0.970 ± 0.021 | 3 | 0.978 ± 0.008 | 0 |
| sz04 | 184 | 0.976 ± 0.047 | 4 | 0.974 ± 0.050 | 12 |
| sz05 | 69 | 0.980 ± 0.011 | 0 | 0.989 ± 0.004 | 0 |
| sz07 | 126 | 0.988 ± 0.010 | 0 | 0.983 ± 0.013 | 0 |
| sz08 | 69 | 0.984 ± 0.007 | 0 | 0.989 ± 0.004 | 0 |
| sz09 | 107 | 0.988 ± 0.005 | 0 | 0.990 ± 0.006 | 0 |
| sz10 | 130 | 0.974 ± 0.047 | 5 | 0.989 ± 0.007 | 0 |
| sz11 | 153 | 0.986 ± 0.005 | 0 | 0.982 ± 0.012 | 0 |
| sz13 | 74 | 0.976 ± 0.012 | 0 | 0.983 ± 0.009 | 0 |
| sz15 | 82 | 0.977 ± 0.015 | 0 | 0.987 ± 0.010 | 0 |
| sz16 | 122 | 0.986 ± 0.007 | 0 | 0.989 ± 0.005 | 0 |
| sz17 | 105 | 0.983 ± 0.009 | 0 | 0.987 ± 0.009 | 0 |
| All | 1525 | 0.980 ± 0.025 | 13 | 0.984 ± 0.020 | 12 |

**Supplementary Table 3. FOOOF goodness-of-fit statistics per participant.** For each participant, the table reports the number of bipolar channels (N), the mean ± standard deviation of the FOOOF $R^2$, and the number of channels with $R^2 < 0.90$, separately for the wakefulness and anesthesia conditions. $R^2$ quantifies the proportion of variance in the log-power spectrum explained by the full FOOOF model (aperiodic + peak components). Across all channel-level fits, model quality was uniformly high (overall mean $R^2 = 0.982$). Only 25 fits (0.8%) fell below $R^2 = 0.90$, and no channels were excluded on this basis.
